# *De novo* biosynthesis of cannabinoid and its analogs in *Yarrowia lipolytica*

**DOI:** 10.1101/2025.02.23.639773

**Authors:** Yuxiang Hong, Yang Gu, Dewei Lin, Zizhao Wu, Wenhao Chen, Tianjian Lu, Pornpatsorn Lertphadungkit, Jingbo Ma, Haili Wang, Bo Zhou, Gil Bar-Sela, Idan Cohen, Peng Xu

## Abstract

*Cannabis sativa* has long been a cornerstone of both medicinal and cultural practices, with its therapeutic use spanning over 2,700 years. Central to its therapeutic effects are cannabinoids, which interact with the endocannabinoid system to influence various physiological processes such as anxiety, pain, and inflammation. Despite its benefits, cannabinoid production faces challenges and scarcity from plant extraction. This work leverages *Yarrowia lipolytica* as a platform for cannabinoid biosynthesis. By optimizing precursor supply, engineering biomolecular condensate-like dual prenyltransferase expression and expanding endogenous metabolism with a noncanonical polyketide synthase, we achieved the *de novo* biosynthesis of various cannabinoid and its analogs. Our engineered *Y. lipolytica* produced ∼3.5 mg/L of cannabigerolic acid, 18.8 mg/L of orsellinic acid, and 0.5 mg/L of cannabigerorcinic acid. Additionally, CBGA titer reached 15.7 mg/L with olivetolic acid supplement. This work demonstrates *Y. lipolytica*’s versatility as a promising host for cannabinoid and its analogs production, which opens avenues for further research and medicinal applications.

## Introduction

*Cannabis sativa* has played a significant role in social and religious practices throughout human history and has been employed therapeutically for over 2,700 years, beginning in ancient China [1–3]. The endocannabinoid system (ECS) is a crucial physiological network implicated in numerous fundamental functions, including neuroplasticity, learning, memory, neuronal development, cell fate determination, nociception, inflammation, appetite, digestion, and the regulation of stress and emotions [4]. Phyto-cannabinoids are a diverse range of chemical compounds known to interact with cannabinoid receptors (CBRs) within the ECS [5, 6]. Cannabidiol (CBD), or CBD-C5, distinguished by its pentyl (five-carbon) side chain, is a major non-psychoactive constituent of *C. sativa*. CBD modulates CBR activation, either enhancing or inhibiting receptor activity, and is implicated in various physiological processes such as anxiety, insomnia, chronic pain, and inflammation [7, 8]. The U.S. Food and Drug Administration (FDA) has acknowledged its therapeutic potential by approving Epidiolex, a purified form of CBD, for the treatment of seizures associated with Lennox-Gastaut syndrome (LGS), Dravet syndrome (DS), and tuberous sclerosis complex (TSC) in patients aged one year and older [9, 10].

Apart from CBD, other CBD analogs have drawn significant attention in medicinal and clinical tests. Cannabigerolic acid (CBGA), another key cannabinoid, acts as the primary precursor for all pentyl side chain cannabinoids. Like CBD, CBGA exhibits anticonvulsant properties in animal models of epilepsy and functions as a dual PPARα/γ agonist, suggesting potential in treating metabolic disorders such as diabetes and dyslipidemia [11]. Notably, CBGA also demonstrated the second-highest inhibitory activity out of all the phytocannabinoids against the COVID-19 3C-like protease [12]. Furthermore, a 2022 preclinical study indicated that CBGA, along with cannabidiolic acid (CBDA) and tetrahydrocannabinolic acid (THCA), could prevent SARS-CoV-2 infection [13]. These findings highlight CBGA’s significant promise for therapeutic development, both as a precursor to other cannabinoids and as a bioactive compound with potential applications in metabolic and antiviral treatments.

Compared to plant extraction, microbial metabolic engineering provides an alternate solution for addressing environmental concerns and resource scarcity [14]. Microbes offer several advantages: they require less arable land, are not affected by climate change, are genetically tractable, and can be scaled up for large-scale production [15, 16]. Additionally, microbes exhibit robust growth and high conversion rates when utilizing a wide range of low-cost renewable raw materials. Nevertheless, only a few groups completed the *de novo* biosynthesis of cannabinoids in microbes. Luo *et al.* first reported the *de novo* biosynthesis of tetrahydrocannabinolic acid (THCA) and cannabidiolic acid (CBDA) in *Saccharomyces cerevisiae*, attaining titers of 8 mg/L THCA and 4.3 μg/L CBDA [17]. Gülck *et al.* attempted cannabinoid biosynthesis using the tobacco plant (*Nicotiana benthamiana*) and *S. cerevisiae*, by feeding OLA as substrate, they got CBGA production at 1 mg/L in the tobacco plant and 5.3 μg/L in yeast [18].

*Yarrowia lipolytica* is a nonconventional yeast and is classified as “generally regarded as safe” (GRAS) [19]. Unlike *S. cerevisiae*, *Y. lipolytica* produces high amounts of acetyl-CoA due to its cytosolic ATP citrate lyase. It also accommodates high flux for malonyl-CoA, and HMG-CoA, and can grow on a wide range of inexpensive raw materials [16, 20]. The oil-accumulating property and expanded inner membrane space facilitate the regio- and stereo-selectivity of various cannabinoids-active enzymes [16, 21]. These attributes make *Y. lipolytica* a superior platform for producing cannabinoids and their analogs.

In this work, we achieved *de novo* biosynthesis of cannabinoid family CBD-C5 and CBD-C1 precursors, including cannabigerolic acid (CBGA), orsellinic acid (OSA), and cannabigerorcinic acid (CBGOA), in *Y. lipolytica*. By combining modular metabolic design, chromosomal targeting knockout, 26_S_ rDNA recombination, and Cre-loxP system, the engineered yeast produced 3.5 mg/L CBGA, 0.5 mg/L CBGOA, and 18.8 mg/L OSA. Additionally, CBGA production reached 15.7 mg/L with olivetolic acid supplement, highest titer reported in the oleaginous yeast culture. This work underscores the potential of *Y. lipolytica* as a versatile chassis for the synthesis of complex terpenes or polyketides. To date, there have been no reports of producing these cannabinoids and their analogues in *Y. lipolytica*. This study not only demonstrates the potential of *Y. lipolytica* as a robust host for complex biosynthetic pathways but also paves the way for future research and medicinal applications. Optimizing production yields and scaling-up the process will be crucial steps toward commercial viability.

## Materials and Methods

### Strains, plasmids, chemicals, reagents and primers

The *Yarrowia lipolytica* strains utilized in this study are outlined in **Supplementary Table S1**. Primers were synthesized by GENEWIZ, Inc. in Guangzhou, China. The specific primers employed in the processes of plasmid assembly, gene sequencing, and colony PCR are comprehensively detailed in **Supplementary Table S2**. Chemicals, including Cannabidiol (CBD-C5, CAS No. 13956-29-1), Cannabidiorocol (CBD-C1, CAS No. 35482-50-9), Cannabigerolic Acid (CBGA, CAS No. 25555-57-1), Cannabigerorcinic Acid (CBGOA, CAS No.69734-83-4), Cannabidiolic Acid (CBDA, CAS No. 1244-58-2), Orsellinic Acid (CAS No. 480-64-8), and Olivetolic Acid (CAS No. 491-72-5), Ro 48-8071 ( Inhibitor of ERG7, CAS No.189197-69-1) were purchased from Cayman Chemical (USA), Shanghai Macklin and Aladdin Biochemical Co., Ltd., Yeast extract powder and yeast peptone were procured from Shanghai Sangon Biotech Co., Ltd., Amino acid including Leucine and uracil were sourced from Shanghai Macklin Biochemical Co., Ltd. For the growth and maintenance of selected yeast strains, we ordered YNB (Yeast Nitrogen Base) as well as two drop-out supplement mixtures: CSM-Ura and CSM-Leu from Sunrise Science, United States. We acquired a plasmid extraction kit and DNA gel purification kits from TIANGEN Biotech, Beijing.

### Recombinant plasmids construction

The genes PpLvaE, CsOLS, CsOAC, CsPT4, NphB, ScERG20, EfmvaE, and EfmvaS were meticulously codon-optimized to enhance their expression in a suitable host. These optimized sequences were procured from Shanghai Sangon Biotech. In addition, the gene ArmB was synthesized specifically for this study by GENEWIZ in Suzhou. For the experimental work, we utilized the YaliBrick plasmid pYLXP’, which has been specifically engineered for facilitating efficient gene expression in the yeast species *Yarrowia lipolytica*. This plasmid offers a versatile framework for cloning and expressing the target genes, making it an ideal tool for our research purposes. Detailed codon-optimized sequences for all these genes can be found in the **Supplementary Sequences** section of our study [22]. To optimize gene expression, we selected three medium-strong promoters and terminators to enable various enzyme expression combinations: TEF-XPR2, P4-PEX20, and pTHD1-MIG1t (**Supplementary Fig. S1**).

We describe two examples of plasmid construction: To create the single-gene expression plasmid pYLXP’-CsOLS, we first linearized the pYLXP’ plasmid using the restriction enzymes SnaBI and KpnI. Next, we integrated a DNA fragment of the CsOLS gene in linearized pYLXP’, previously amplified by PCR, through Gibson Assembly [23]. Finally, we verified the accuracy of the pYLXP’-CsOLS plasmid via sequencing conducted by experts at GENEWIZ, Inc. in Guangzhou, China, ensuring it was ready for the next stage of research.

To create the multi-gene expression plasmid pYLXP’-CsOAC-CsOLS, we employed the subcloning method. We initiated the process by linearizing the pYLXP’-CsOAC plasmid using the restriction enzymes NheI and SalI. This step was crucial as it created specific ends for subsequent ligation. Concurrently, we prepared the CsOLS expression cassette by digesting the pYLXP’-CsOLS plasmid with the restriction enzymes AvrII and SalI. This digestion yielded fragments that included the necessary coding sequence for CsOLS. Following the digestion steps, we proceeded to ligate the linearized pYLXP’-CsOAC with the CsOLS expression cassette using T4 DNA ligase allowing us to construct the recombinant pYLXP’-CsOAC-CsOLS plasmid. To ensure the successful formation of this new plasmid, we conducted verification through endonuclease digestion, confirming that the expected banding pattern was present on an agarose gel.

### Gene knockout in Yarrowia lipolytica

We employed Cre-loxP recombination system to delete specific genes in *Yarrowia lipolytica* [20], aiming for gene knockout or integration at chromosome X loci (see **Supplementary Fig. S2**). For this purpose, we employed the integrative plasmid pUrlp-X-loci [20], which is based on the pYLXP’ mentioned above.

This example outlines the knockout process for the DGA1 gene. Initially, we PCR-amplified the upstream and downstream sequences surrounding the DGA1 gene, designated as DGA1-Up and DGA1-Dw. We then digested the plasmid pUrlp using the restriction enzymes AvrII and SalI, which produced a loxP-Ura-loxP cassette and a linearized version of pUrlp. This linearized plasmid was then purified through gel electrophoresis. Following purification, we employed Gibson Assembly to combine the following fragments: the linearized pUrlp, DGA1-Up, the loxP-Ura-loxP cassette, and DGA1-Dw. This assembly yielded the knockout plasmid pUrlp-ΔDGA1, which was sequenced by GENEWIZcin Guangzhou

Once sequencing was complete, we extracted the knockout cassette for the DGA1 gene from pUrlp-ΔDGA1 using endonuclease digestion and introduced it into *Y. lipolytica* through transformation. We identified positive transformants through colony PCR. After this, we introduced the plasmid pYLXP’-Cre, which encodes Cre recombinase, to remove the auxotrophic marker Ura3 from these transformants. To eliminate pYLXP’-Cre plasmid from the modified *Yarrowia lipolytica*, we incubated the cells in rich YPD liquid medium at 28-30 °C for a duration of 24 to 48 hours.

### Genomic integration in targeted loci of the engineered pathway

To maintain genetic stability, a variety of promoters and terminators were employed to control the expression of multiple genes. This strategy involved three strong promoters (pFBA1, pTDH1, and pTEF) paired with three terminators (PEX20, MIG1, and XPR2). In this study, we selected the targeted loci integration YALI0A08734g, YALI0C08701g (Ku70), YALI0C01221g, YALI0C06446g(DLD2), YALI0D15246g, YALI0E27654g(POX4), YALI0E20977g(ARO8), YALI0D07986g(DGA2), YALI0E32769g (DGA1), YALI0D00789g(GGPPS), YALI0E03212g(LDH) and YALI0F26169g, detail was described in previous report (**Supplementary Fig. S2**) [20]. The process of integrating the multiple gene cassettes into genomic loci resembles targeted gene knockout.

### Yeast transformation, sample preparation and extraction

The Li-Ac protocol for transforming *Y. lipolytica* has been effectively documented in studies by Ma et al., showcasing its reliability and efficiency in genetic modifications [21]. For product extraction, 400 µL of whole cell culture was collected and processed using a 3D centrifugal cryogenic sample grinder (Jingxin JXCL-6K, Shanghai, China) with 400 µL ethyl acetate (0.05% formic acid) and glass beads (0.5 mm) at −20 °C. The grinding was performed at 21 m/s for 40 seconds with a 20-second interval, repeated for 40 cycles. After the initial extraction, 300 µL of the organic layer was transferred to a fresh microfuge tube. Two additional extractions were performed, each with 400 µL ethyl acetate (0.05% formic acid), and 300 µL of the upper organic layer was added to the previous extracts. The organic fraction was then evaporated using an Eppendorf Speedvac Concentrator, and the dried extracts were resuspended in methanol/HLO (80/20) and vortexed for 1 minute [21].

### HPLC and LC-MS quantification of cannabinoids products

The samples underwent analysis using high-performance liquid chromatography (HPLC) with a 1260 Infinity liquid chromatograph. The system utilized a reverse-phase C18 column (ZORBAX Eclipse Plus C18, 4.6×100 mm with a particle size of 3.5 microns, Agilent Technologies). For detection, a diode array detector was employed, calibrated to wavelengths of 210 nm and 270 nm to capture the necessary absorbance of the analyzed compounds.

In terms of the mobile phase, it consisted of two distinct solvents: solvent A was prepared with purified water supplemented with 0.05% formic acid, serving primarily as the aqueous component, while solvent B was comprised of high-purity methanol. The separation of various compounds was achieved through a carefully controlled gradient elution process. Initially, the ratio of solvent B was set at 60%, gradually increased to 100% over a span of 20 minutes to enhance the separation of target analytes. This high concentration of solvent B was maintained for 4 minutes to ensure complete elution of strongly retained components. Subsequently, the solvent composition was adjusted, decreasing from 100% back to 60% over the course of 1 minute, followed by a stabilization period of 2 minutes at 60% solvent B to reset the system for the next analysis cycle.

The LC-MS analysis was conducted using a Thermo Scientific Orbitrap Exploris 120 mass spectrometer equipped with a heatable electrospray ionization (HESI) source. Chromatographic separation was performed on an Accucore C18 HPLC column (2.6 µm, 2.1 x 150 mm). The mobile phase utilized in the experiment consisted of two components: 0.1% formic acid dissolved in water, which is referred to as phase A, and methanol, identified as phase B. The gradient elution was meticulously programmed as follows: at the start (0 minutes), the composition was set to 5% phase B. This initial concentration was maintained for the first half-minute (0.5 minutes) to establish a stable baseline. As the run progressed, the proportion of phase B increased to 95% at 13 minutes, allowing for robust separation of the analytes. This high concentration of methanol was held constant until 16.9 minutes before the timing shifted again. At 17 minutes, the mobile phase composition rapidly returned to the initial 5% phase B, effectively regenerating the column for the next run. The flow rate was controlled at 0.3 mL/min, and each injection during the analysis was carefully measured at 1 µL. Mass spectrometric detection was carried out in the mass range of 100 to 1000 m/z, employing collision energies of 20, 50, and 80 electron volts (eV) to facilitate the fragmentation of ions and enhance the specificity of detection. The identification of compounds was achieved through an automated search process that utilized a variety of comprehensive databases and spectral libraries. Notable online database resources included mzCloud, KEGG, ChemSpider, mzVault spectral library, and BioCyc.

### Confocal laser scanning microscope

Imaging was conducted using a Zeiss LSM 980 confocal laser scanning microscope equipped with Airyscan 2 technology. Yeast cells were prepared and mounted on glass slides for optimal visualization. The Airyscan 2 detector allowed for super-resolution imaging, enhancing the spatial resolution beyond the diffraction limit of light. This was achieved by collecting more spatial information through a unique detector design, resulting in high-definition images with an improved signal-to-noise ratio. The system was calibrated and optimized for various fluorophores, including Nile red, TurboGFP and mScarlet channels, to ensure accurate and reproducible results. Z-stack imaging was performed to capture three-dimensional structures, and image processing was carried out using Zeiss ZEN software, enabling detailed analysis and visualization of subcellular components labeled with turboGFP and mScarlet or stained with Nile red.

### Statistics

Experiments were independently repeated three times to ensure reliable results, presented with their standard deviations (SD). A two-sided t-test was used for statistical analysis, with p-values calculated using Origin 2024 or GraphPad Prism 10.1.

## Results and Discussion

### Optimizing hexanoyl-CoA and malonyl-CoA supply to improve CBD-C5 precursor olivetolic acid

The biosynthetic pathway for CBD-C5 involves three modules: the polyketide olivetolic acid (OLA) pathway, the isoprenoid pathway producing geranyl pyrophosphate (GPP), and a prenyltransfer step synthesizing cannabigerolic acid (CBGA), the precursor to various cannabinoids (Fig. 1a). OLA is derived from one hexanoyl-CoA and three malonyl-CoA units, catalyzed by olivetol synthase (OLS), which extends one unit of hexanoyl-CoA with three units of malonyl-CoA. Then the tetraketide intermediate cyclizes to form OLA by olivetolic acid cyclase (OAC). Supply of hexanoyl-CoA is the rate-limiting step for CBD synthesis [21]. CsAAE1 from *Cannabis sativa* (C. sativa) has been successfully expressed in yeast to achieve a trace amount of CBDs [17]. Our group previously identified the LvaE gene encoding the short-chain acyl-CoA synthetase from *Pseudomonas putida* KT2440 (PpLvaE) [21], which efficiently converts hexanoic acid to hexanoyl-CoA, and improved OLA production.

**Figure 1.**
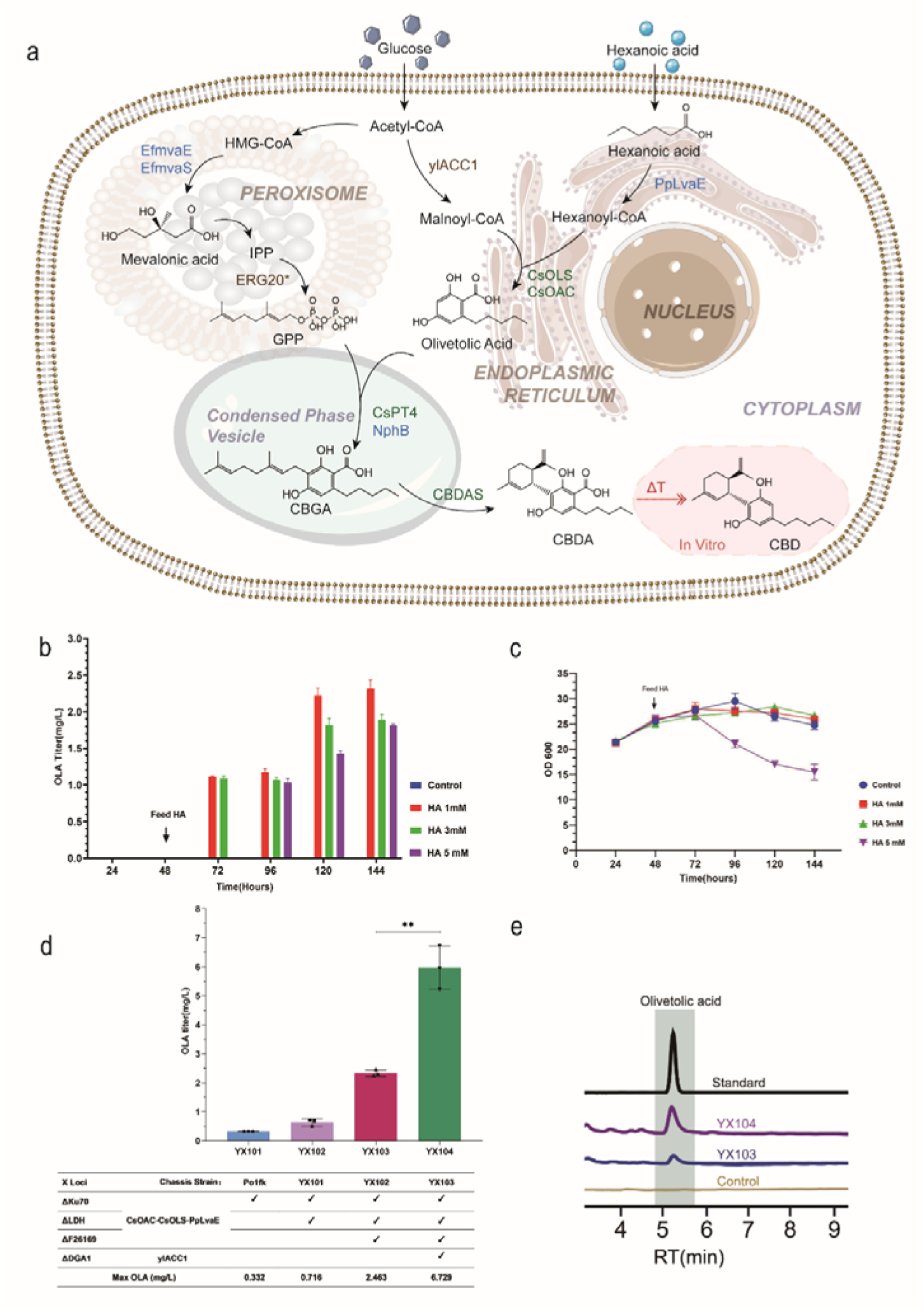
| Biosynthesis of Cannabidiol (CBD-C5) precursor olivetolic acid. (***a***) Cannabinoids biosynthesis pathway in *Y. lipolytica*, (***b*)**. Olivetolic acid titer in YX103 with 1 mM, 3 mM, and 5 mM hexanoic acid (HA) supplementation at 48 hours, (***c***). OD600 of YX103 with 1 mM, 3 mM, and 5 mM hexanoic acid (HA) supplementation at 48 hours, (***d***). Relevant gene expression characteristics and OLA titer of YX101,YX102, YX103, and YX104. (***e***). HPLC results of olivetolic acid in strains. Values are shown as mean ± SD (*n* = 3 biologically independent replicates). Two-tailed Welch’s t-test was used to compare two groups and *p* values are shown.

Hexanoic acid feeding above 1 mM was found to be detrimental to cell growth (Fig. 1b and 1c). Integration of the OLA cassette into different chromosomal loci significantly improved OLA production. Strain YX101, with the OLA cassette integrated at the YALI0C08701g (Ku70) locus, produced 0.33 mg/L of OLA with 1 mM hexanoic acid (HA) supplementation (Fig.1b). Subsequent integration of a second OLA cassette at YALI0E03212g (LDH) in strain YX102 doubled OLA production to 0.72 mg/L (Fig. 1d). Further integration at the YALI0F26169g locus (ATP-dependent Lon protease) in strain YX103, combined with delayed HA supplementation (48 hours post-inoculation) at 1 mM, increased production to 2.46 mg/L at 144 hours (Fig. 1d,1e). OLA production was confirmed by HPLC and high-resolution LC-MS (Figure 1e, **Supplementary Fig. S3**).

To mitigate acetyl-CoA flowing to the lipid pathway and enhance malonyl-CoA availability, we further knocked out DGA1 (diglycerol acyltransferase) and overexpressed ylACC1 (acetyl-CoA carboxylase), resulting in strain YX104 achieving an OLA titer of 6.73 mg/L, a threefold increase over YX103 (Figure 1d,1e). This result indicates that diverting acetyl-CoA flux from TAG synthesis is effective to improve OLA level in *Y. lipolytica*.

### Optimizing GPP supply and prenyltransferase module to improve CBGA production

Geranyl diphosphate (GPP) supply is a rate-limiting factor in terpene biosynthesis (Figure 1). The cytosolic farnesyl pyrophosphate synthase (ERG20) acts as a bifunctional catalyst, sequentially condensing isopentenyl diphosphate (IPP) and dimethylallyl pyrophosphate (DMAPP), to form farnesyl diphosphate (FPP). Previously, Codruta *et al.* utilized a mutant version of ScERG20\**^F96W/N127W^* from *S. cerevisiae* that preferentially produces GPP over FPP and achieved a significant increase in monoterpene titers [24]. Herein we used both mutant ylERG20\**^F88W/N119W^* from *Y. lipolytica* and ScERG20\**^F96W/N127W^* from *S. cerevisiae* which are supposed to perform preferential GPP production (Fig. 1a).

Besides, to boost IPP and DMAPP precursors, we also leveraged heterologous genes *mvaE* and *mvaS* from *Enterococcus faecalis* to replace the native ERG10, ERG13, and tHMGR (Figure 1) [25]. These enzymes catalyze the first three steps of the MVA pathway and have proven high efficiency in *S. cerevisiae* [25]. We integrated EfmvaE, EfmvaS, ylERG20\**^F88W/N119W^* at the YALI0A08734g locus, while *ScERG*20\**^F96W/N127W^* at geranylgeranyl pyrophosphate synthase (GGPPS) locus, and these strategies increased GPP flux in the peroxisome. Combined with prenyltransferase (PTase) integration, we will evaluate the strains’ performance on CBGA production (Fig.1a, 3a, and 3d).

Prenyltransferase (PTase) is another rate-limiting step in CBGA pathway (Fig. 1a). The PTase from *Cannabis sativa* (CsPT4) has been reported to play a crucial role in CBGA biosynthesis. The first 77 amino acids coding for chloroplast-targeting signal has been removed for better activity [17]. With this, we engineered a truncated CsPT4 devoid of the first 77 amino acids named tCsPT4. Plant enzymes with truncated chloroplast-targeting sequences may present in the cytoplasm or endoplasmic reticulum (ER) of yeast cells. To determine the exact cellular localization of tCsPT4 in *Y. lipolytica*, we applied confocal microscopy to identify its spatial distribution.

As an ER-positive control, we fused TurboGFP to the C-terminal of two ER retention signals: ER1 or ER2 (**Supplementary Fig. S4**). These tags are known to localize enzymes to the ER lumen or the cytoplasmic face of the ER membrane in *S. cerevisiae* [26]. Under confocal fluorescence microscopy, our positive control TurboGFP-ER1/ER2 is successfully localized to the intracellular compartment, including the endoplasmic reticulum (Fig. 2a). For tCsPT4, we fused its C-terminal with mScarlet and co-expressed TurboGFP-ER1/ER2 and tCsPT4-mScarlet in the same strain (Figure 2b, **Supplementary Fig. S4**). Merged confocal images showed that tCsPT4 was predominantly localized onto the expanded ER surface (Fig. 2b).

**Figure 2.**
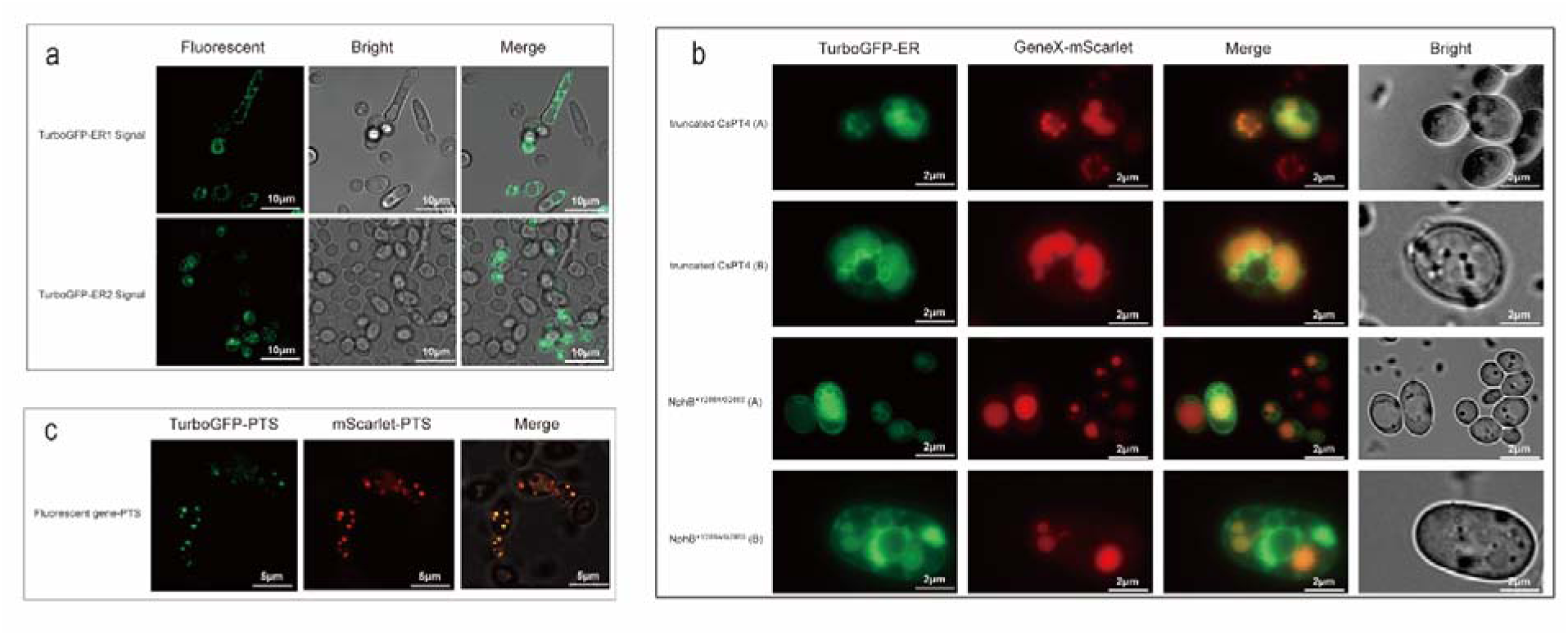
| Prenyltransferase Confocal fluorescence microscopy verification and biosynthesis of CBGA. (***a***). Confocal fluorescence microscopy image of TurboGFP fused ER1/ER2 expression in *Y. lipolytica*, (***b***). Confocal fluorescence microscopy image of tCSPT4 and *NphB*^Y288A/G286S^*expression in *Y. lipolytica*, (***c***). Confocal fluorescence microscopy image of PTS signal tag expression in *Y. lipolytica*.

Peroxisomal β-oxidation pathway can directly supply acetyl-CoA from fatty acids degradation. The peroxisomal localization of specialized plant pathways has previously enabled efficient squalene and alkaloid production [27, 28]. We next attempted localizing tCsPT4 to peroxisomes. We have validated that the SKL signal successfully directed proteins to the peroxisomes of *Y. lipolytica* (Fig. 2C).

We fused SKL with tCsPT4 and integrated this engineered PT4 at YALI0C01221g (encoding 26s proteasome non-ATPase regulatory subunit 10), resulting in strains YX105(Without PTS) and YX106 (With PTS) (Fig. 3a). Strain YX105, without the PTS signal, produced 7.77 mg/L OLA and 143 µg/L CBGA (Fig. 3a). In contrast, YX106 with PTS signal, produces 5.56 mg/L OLA and without detectable CBGA (Fig. 3a). This suggests that PTase peroxisome targeting is not suitable to produce OLA or CBGA. Next, we enhanced GPP flux in YX105 by integrating Ef*mvaE*, Ef*mvaS*, and ylERG20\**^F88W/N119W^*. The engineered YX107 produced 3.39 mg/L of OLA and 335 µg/L of CBGA (Fig. 3a, **Supplementary Fig. S5**), representing a 2.34-fold increase compared to YX105.

**Figure 3.**
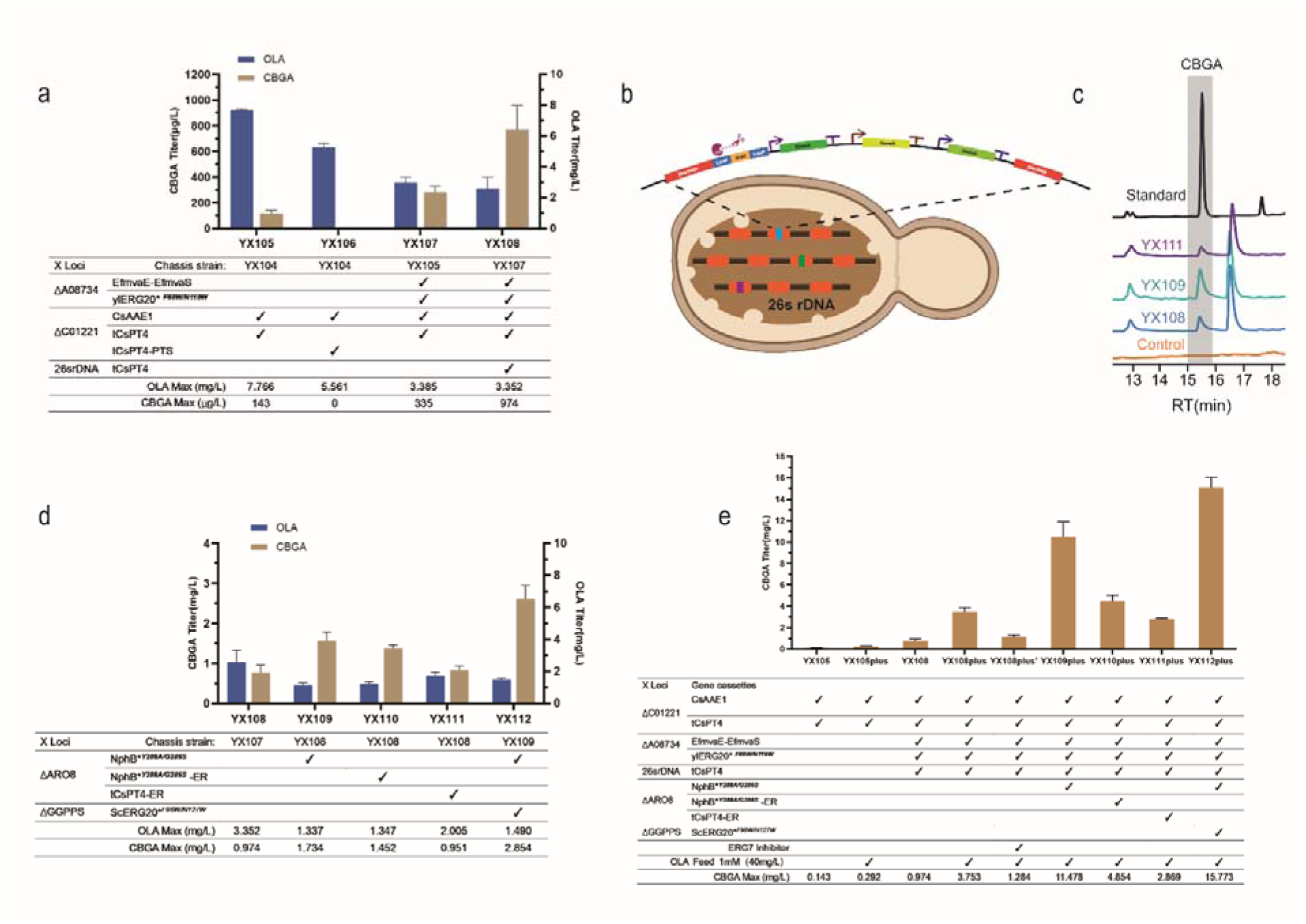
| Prenyltransferase optimization and biosynthesis of CBGA. (***a***). Relevant gene expression characteristics; OLA and CBGA titer of YX105, YX106, YX107, and YX108, (***b*)**. 26S rDNA recombination and the Cre-loxP System strategy scheme, (***c***). HPLC results of CBGA, (***d***). Relevant gene expression characteristics; OLA and CBGA titer of YX108, YX109, YX111, and YX112, (***e***). Strains YX108plus, YX109plus, YX110plus, YX111plus, and YX112plus and their CBGA titer supplemented additional 1mM OLA.

As PTase remained a bottleneck, we hypothesized that increasing the copy number of tCsPT4 would enhance CBGA production. *Y. lipolytica* contains at least 200 copies of ribosomal DNA (rDNA) clusters [29]. These rDNAs can be leveraged for random integration of heterologous pathways (Fig. 3b) [30]. In order to pull away the GPP flux toward CBGA synthesis, we increased the copy number of tCsPT4 with the 26s rDNA and Cre-loxP System (Fig. 3b). The engineered strain YX108, produced 3.35 mg/L OLA and 974 µg/L CBGA (Fig. 3a,3b,3c), representing a 3-fold CBGA increase compared to YX107.

### Condensed phase vesicle-like dual PTase expression improved CBGA production

In strain YX108, despite increasing the GPP flux and the copy numbers of tCsPT4, CBGA production remained below 1 mg/L. We hypothesized that low CBGA production might be due to suboptimal catalytic activity of tCsPT4 or pathway incompatibility with the host organism. To address this challenge, we chose to co-express a different PTase, NphB from *Streptomyces sp.*, which offers complementary prenyltransferase activity, to improve CBGA production [31–33]. NphB is a soluble aromatic PTase initially characterized by *Kuzuyama et al.* [34]. Studies on mutant versions of NphB, specifically *Y288A* and *G286S*, have shown improved catalytic specificity [32]. We analyzed the subcellular distribution of NphB*\*^Y288A/G286S^* in *Y. lipolytica* with confocal fluorescence microscopy and confirmed NphB*\*^Y288A/G286S^* localized into expanded ER-like cell vesicles, similar to tCsPT4 (Fig. 2b). Then we integrated NphB\**^Y288A/G286S^* into the chromosomal locus YALI0E20977g (ARO8) of YX108, generating strain YX109. And this led to the production of 1.73 mg/L of CBGA, which is almost 2-fold increase compared to YX108 (Fig. 3d).

As the endoplasmic reticulum (ER) is the location where OLA synthesis takes place (Fig.1a), Zhang *et al.* showed that localizing PTase onto the ER surface can improve CBGA production in *S. cerevisiae* [35]. To improve the catalytic efficiency of tCsPT4 and NphB, we fused an ER signal tag (validated previously) to the N-terminus of both enzymes. We integrated NphB\**^Y288A/G286S^*-ER, or tCsPT4-ER into the chromosomal locus ARO8 of YX108, generating strains YX110, and YX111. And YX110 produced 1.45 mg/L of CBGA, while YX111 produced 0.93 mg/L of CBGA (Fig. 3d), indicating ER targeting is not benefiting CBGA synthesis.

Notably, YX109, which expresses NphB\**^Y288A/G286S^* without the ER tag, achieved the highest CBGA production (1.73 mg/L), representing a 12-fold increase compared to the original YX105 strain (Fig. 3d). Furthermore, after increasing the GPP flux in YX109 by replacing endogenous GGPPS with *ScERG*20\**^F96W/N127W^*, we obtained strain YX112, which produces 2.85 mg/L CBGA, the highest titer compared to the previous strains (Fig. 3d).

Due to the fact that both PTases, tCsPT4 and NphB\**^Y288A/G286S^*, are spatially assembled together forming a condensed phase vesicle-like structure (Fig. 2b), we assume that dual PTase co-expression may be similar to liquid-liquid phase separation induced biomolecular condensates [36]. This condensed vesicle-like PTase expression significantly enhances CBGA production (2.85 mg/L) in *Y. lipolytica*. While OLA precursor remains a bottleneck in the chassis strain (OLA titer less than 10 mg/L), we next probed the potential of the condensate-like PTase, by feeding 1 mM (∼40 mg/L) of OLA to the strains YX108, YX109, YX110, YX111 and YX112 (Fig. 3e). Furthermore, we applied an ERG7 inhibitor (Ro 48-8071 fumarate which inhibits oxidosqualene cyclase) to boost GPP precursor [37]. Surprisingly, strains YX109plus and YX112plus with condensed phase vesicle-like dual PTase expression, achieved the highest CBGA production (11.48 mg/L and 15.77 mg/L) (Fig. 3e). Notably, YX112 showed the highest increase, producing over 15.77 mg/L of CBGA with about 39% conversion ratiofrom OLA to CBGA (Fig. 3e). Machine-learning based protein sequence analysis [38] indicates that tCsPT4 and NphB will form a biomolecular condensate at a very high probability.

This result demonstrates condensed phase vesicle-like PTase expression, significantly enhanced CBGA production in *Y. lipolytica*, which also reinforced that OLA precursor is currently the rate-limiting step for further improving CBGA yield in *Y. lipolytica*.

### Biosynthesis of cannabidiorcol precursor orsellinic acid

Cannabidiorcol (CBD-C1 or O-1821) is an underexplored minor cannabinoid derived from *C. sativa*, distinguished by a shortened methyl sidechain (C1) instead of the pentyl group seen in CBD-C5. Like CBD-C5, it is non-psychoactive. However, CBD-C1 remains less studied than other cannabinoids like THC and CBD-C5, although its unique structure offers potential for therapeutic applications [39]. Cannabigerorcinic acid (CBGOA) serves as the precursor for methyl-chain cannabinoids like CBD-C1, much like CBGA does for pentyl cannabinoids [40, 41]. CBGOA’s precursor, orsellinic acid (OSA), is a dihydroxybenzoic acid with antioxidant and neuroprotective properties, commonly found in lichens as a fungal metabolite [41, 42]. Therefore, engineering heterologous production of CBD-C1 analogs and its precursor OSA holds promise to overcome the supply limitations and expand their medicinal applications.

OSA synthesis does not require hexanoyl-CoA, making the process faster and more efficient compared to the OLA pathway (Fig. 4a). We identified an iterative type I polyketide synthase (PKS), orsellinic acid synthase (ArmB), from *Armillaria mellea* [43]. To date, there have been no reports of ArmB being used for heterologous expressions in yeast or *E. coli*. ArmB contains multiple domains, including ACP transacylase (SAT), ketosynthase (KS), malonyl-CoA transacylase (MAT), product template domain (PT), acyl-carrier protein (ACP) and thioesterase domain (TE) (Fig. 4b), with MW around 238 kDa [44, 45].

**Figure 4.**
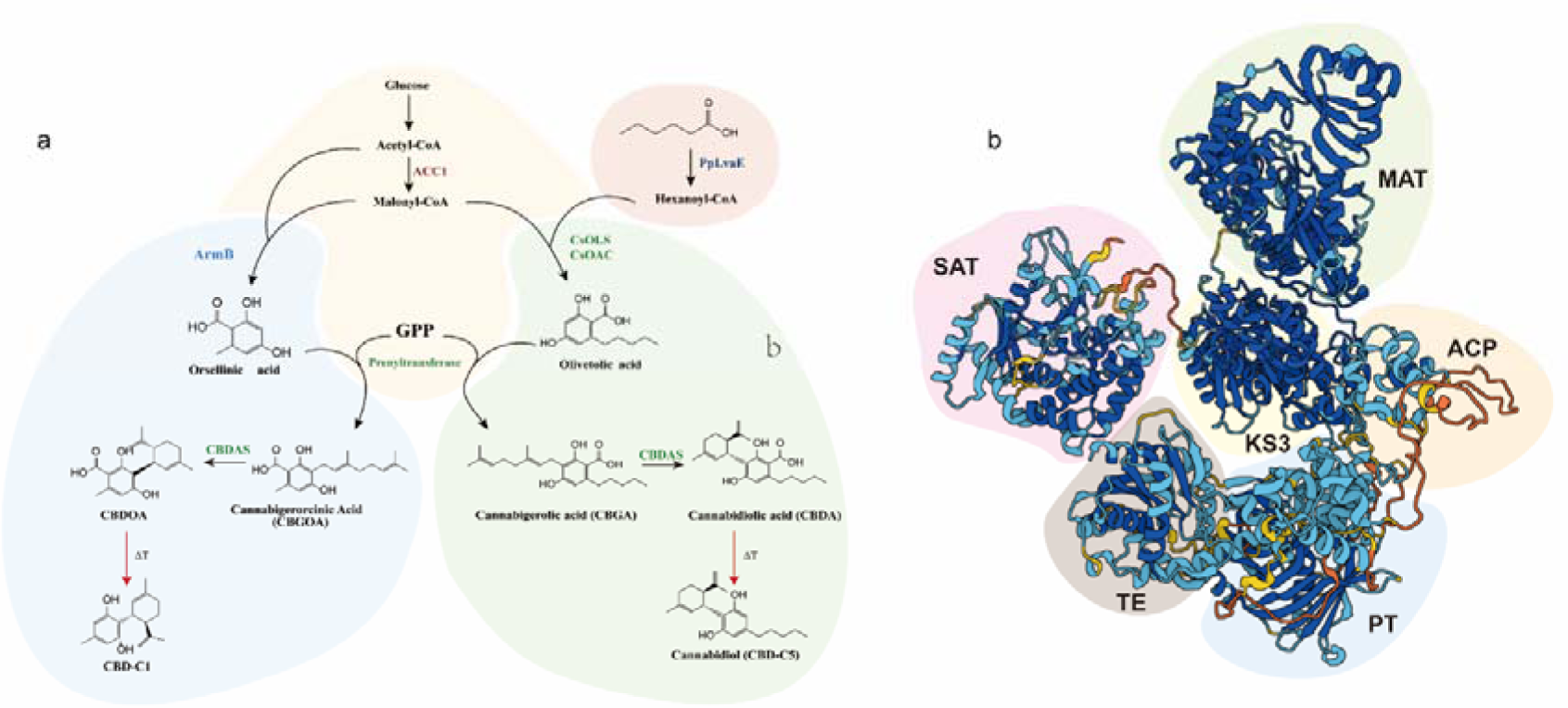
|Biosynthesis pathway of Cannabidiorcol (CBD-C1) (***a***) CBD-C1 biosynthesis pathway compared to CBD-C5, (***b***). Schematic diagram of orsellinic acid synthase (ArmB) protein structure and functional domains.

We next integrated this codon-optimized ArmB into the Ku70 locus in *Y. lipolytica*. Strain YX201 produced 2.19 mg/L of OSA, which was validated by HPLC and high-resolution LC-MS (Fig. 5a,5b, **Supplementary Fig. S6**).

**Figure 5.**
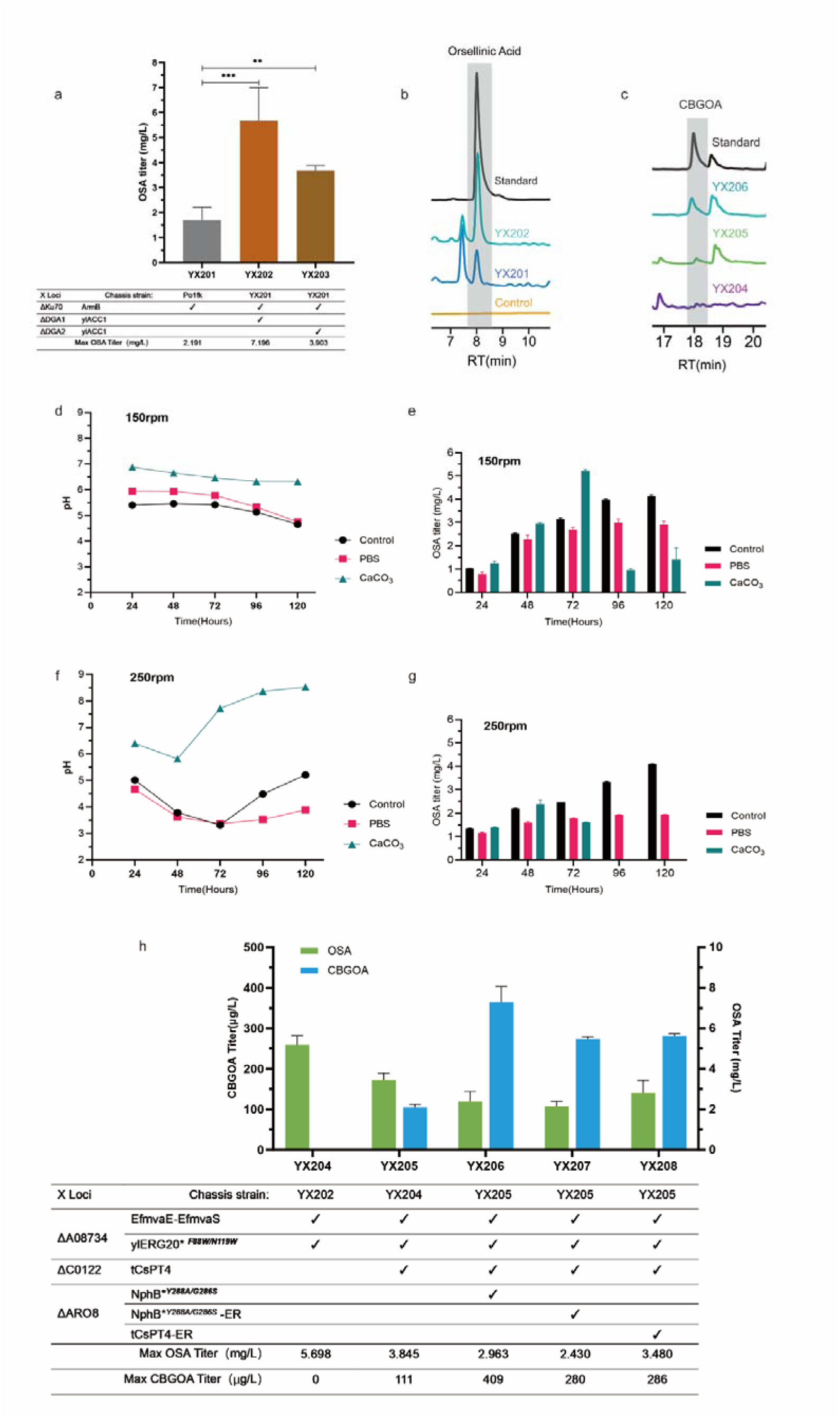
| Biosynthesis of Orsellinic acid and CBGOA. (***a***). Relevant gene expression characteristics and OSA titer of YX201, YX202, and YX203, (***b*)**. HPLC results of OSA, (***c***). HPLC results of CBGOA, (***d***). 5-days fermentation pH values change of YX202 using either PBS buffer or CaCOL with 150rpm, (***e***). 5-day fermentation OSA titer change of YX202 using either PBS buffer or CaCOL with 150rpm, (***f***). 5-days fermentation pH values change of YX202 using either PBS buffer or CaCOL with 250rpm, (***g***). 5-day fermentation OSA titer change of YX202 using either PBS buffer or CaCOL with 250rpm, (***h***). Relevant gene expression characteristics; OLA and CBGA titer of YX204, YX205, YX206, YX207, and YX208. Values are shown as mean ± SD (*n* = 3 biologically independent replicates).

Since OSA synthesis follows Claisen condensation with three malonyl-CoA as the extending units, increasing malonyl-CoA levels is crucial. We integrated *ylACC1* gene cassettes into either the *DGA1* or *DGA2* loci of YX201 to enhance malonyl-CoA supply, creating strains YX202 and YX203. In test tubes, YX202 produced 7.19 mg/L OSA, and YX203 produced 3.90 mg/L (Fig. 5a,5b), a more than 3-fold increase compared to strain YX201.

In the shake flask, we observed culture pH dropped to 4, potentially impairing membrane permeability and strain performance. To address this, we applied PBS buffer or CaCOL and adjusted the agitation rates when cultivating YX202. CaCOL effectively stabilized pH, and OSA titer peaked at 5.278 mg/L at 72 h and 150 rpm. However, OSA began to degrade after 72 h in the presence of CaCOL (Fig. 5d-5g), possibly due to endogenous enzyme activity. PBS was ineffective at both 150 rpm and 250 rpm. The YPD control group showed steady OSA production, exceeding 4 mg/L, despite pH fluctuations. Agitation at 150 rpm was more conducive to strain growth. The strain in YPD medium produced 4.23 mg/L of OSA at 150 rpm (Fig. 5d-5g). This result indicates that CBD-C1 precursor OSA gene cluster can be functionally reconstituted in *Y. lipolytica*.

### Production of CBD-C1 analog cannabigerorcinic acid (CBGOA)

The downstream module of the CBD-C1 biosynthesis pathway closely resembles that of CBD-C5, with PTase transferring a GPP moiety to the OSA side chain (Fig. 4a). To facilitate CBGOA production, we employed the same strategy used for CBGA production. Specifically, we integrated EfmvaE, EfmvaS, and ylERG20\**^F88W/N119W^* into the YALI0A08734g locus of strain YX202, resulting in strain YX204 (Fig. 5c,5h, **Supplementary Fig. S7**).

Single and dual PTase expression were also performed in YX204. Similar to the CBGA result, the highest CBGOA production (409 µg/L) was observed in YX206 which was engineered to harbor a condensate-like dual PTase expression system (Fig. 2b, 5c, and 5h). Dual PTase was less effective for CBGOA than for CBGA, despite using the same enzymes. This discrepancy is likely due to the limited supply of OSA and the subtle structural difference in the substrate, as variations in the side chain geometry and length alter the position of the substrate-GPP alignment within the catalytic pocket, thereby the catalytic efficiency shifted. Thus, further AI-assisted engineering of the PTase or mining of novel PTase that prefers OSA as a substrate is the key to enhance its performance and deliver more potent CBD-C1 analogs.

### Engineering YX112 and YX206 strains Fed-batch cultivation in shake flasks

To optimize the production of YX112 and YX206 engineering strains, we conducted fed-batch cultivation using flat shake flasks (Fig 6a, 6b). For strain YX112, the medium was supplemented with 1 mM hexanoic acid on Day 1. The OD600 exhibited steady growth, reaching approximately 52 by Day 6. Olivetolic acid concentration increased continuously, peaking on Day 4 at approximately 9.84 mg/L, after which it decreased due to its conversion to CBGA. By Day 6, the olivetolic acid titer stabilized at 7.64 mg/L. CBGA production began accumulating on Day 2, with a final titer of approximately 3.537 mg/L by Day 6 (Fig 6a).

**Figure 6.**
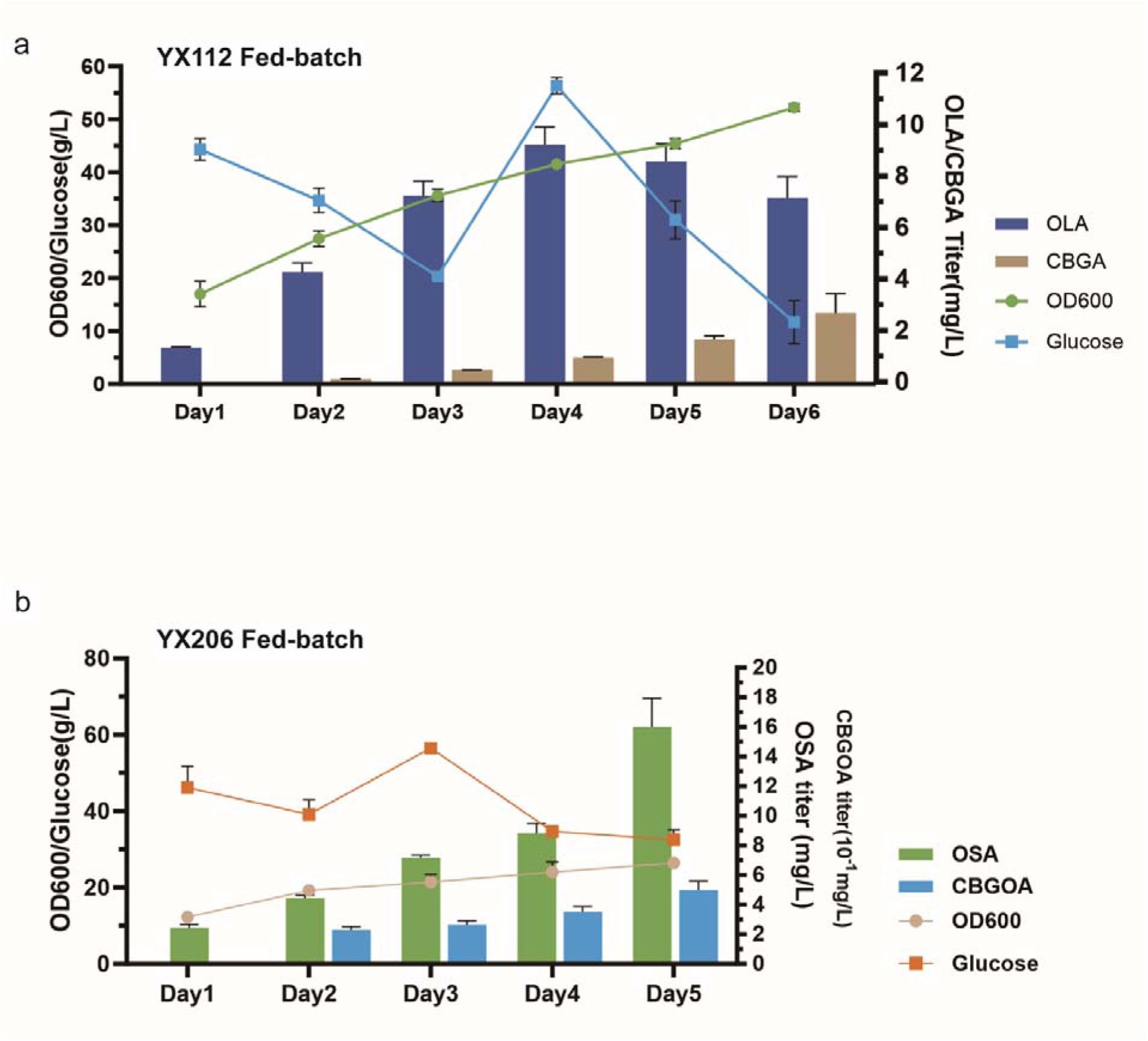
| Fed-batch cultivation in shake flasks. (***a***). YX112 Fed-batch cultivation in shake flasks for 6 days, 1mM hexanoic acid supplementation at Day 1, OD600, Glucose content, OLA, and CBGA were measured every 24 hours, (***b*)**. YX206 Fed-batch cultivation in shake flasks for 5 days, OD600, Glucose content, OSA, and CBGOA were measured every 24 hours. Values are shown as mean ± SD (*n* = 3 biologically independent replicates).

For strain YX206, glucose served as the sole substrate for orsellinic acid (OSA) and CBGOA production. The glucose concentration was maintained at no less than 30 g/L throughout the cultivation period. Although the OD600 increased steadily, it progressed at a slower rate than YX112, reaching approximately 28 by Day 5. Orsellinic acid production continuously increased, with a final titer of 18.87 mg/L. However, CBGOA production was less efficient compared to CBGA, resulting in a final titer of approximately 541 µg/L by Day 5 (Fig 6b).

## Conclusions

In this study, we successfully engineered nonconventional oleaginous yeast *Y. lipolytica* for the *De novo* biosynthesis of CBD and its analogs, including olivetolic acid (OLA), cannabigerolic acid (CBGA), orsellinic acid (OSA), and cannabigerorcinic acid (CBGOA). By optimizing the polyketide pathway for olivetolic acid (OLA) production and boosting geranyl pyrophosphate (GPP) precursor, we overcame the pathway bottlenecks and achieved successful microbial synthesis of cannabinoids. The strategic engineering of prenyltransferases, in particular, we observed a condensate-like expression of dual PTase in the engineered strain, significantly improved CBGA titer to 15.7 mg/L with OLA supplementation.

Our work represents the first reported production of these cannabinoids and their analogs in *Y. lipolytica*, highlighting the yeast’s potential as a versatile chassis for the synthesis of complex terpenes and polyketides. The remarkable titer improvement of CBD analogs demonstrates the effectiveness of our metabolic engineering strategies. These findings not only demonstrate the feasibility of producing medically relevant cannabinoids in a microbial host but also lay the groundwork for future efforts to optimize production yields and scalability. Addressing remaining challenges—such as enhancing OLA availability, improving PTase efficiency, and fully unraveling the enzyme localization and function underlying the CBD pathway — will be crucial for advancing toward commercial viability.

Our study contributes to the development of sustainable and scalable methods for cannabinoid production in *Y. lipolytica*, with potential applications in pharmaceuticals and therapeutics. The approaches described here can also be extended to the biosynthesis of other complex natural products, paving the way for innovative applications in biotechnology and medicine.

## Acknowledgments

We thank the Core Facility team of the Department of Biotechnology and Food Engineers, Guangdong-Israel Institute of Technology for their service and technical support. The funding is supported by the National Natural Science Foundation of China (general grant 22378083) and Li Ka-shing Foundation (EN2400022) and Muyuan Laboratory (Program ID: 12106022401).

## Author contributions

YX.H. designed, and performed experiments, collected and analyzed the data, generated the figures, and wrote the manuscript. Y.G. designed, conducted plasmid construction, and revised the manuscript. JB.M., designed and conducted plasmid construction, DW.L., ZZ.W., WH.C., TJ.L., and P.L. conducted auxiliary experiments, HL.W., B.Z., conducted LC-MS analysis, BSG and CI supervised the work, and P.X., designed, supervised, and sponsored the study and revised the manuscript.

## Data availability

All data from the current study is available from the corresponding author upon request.

## Material availability

All data from the current study is available from the corresponding author upon request.

