## Supplementary Material for "*De novo* biosynthesis of cannabinoid and its analogs in *Yarrowia lipolytica*"

**Supplementary Figure S1 Triple copy of the olivetolic acid pathway integration strategy scheme in this study, we selected three medium-strong promoters and terminators to enable various enzyme expression combinations: TEF-XPR2, P4-PEX20, and pTHD1-MIG1t.**

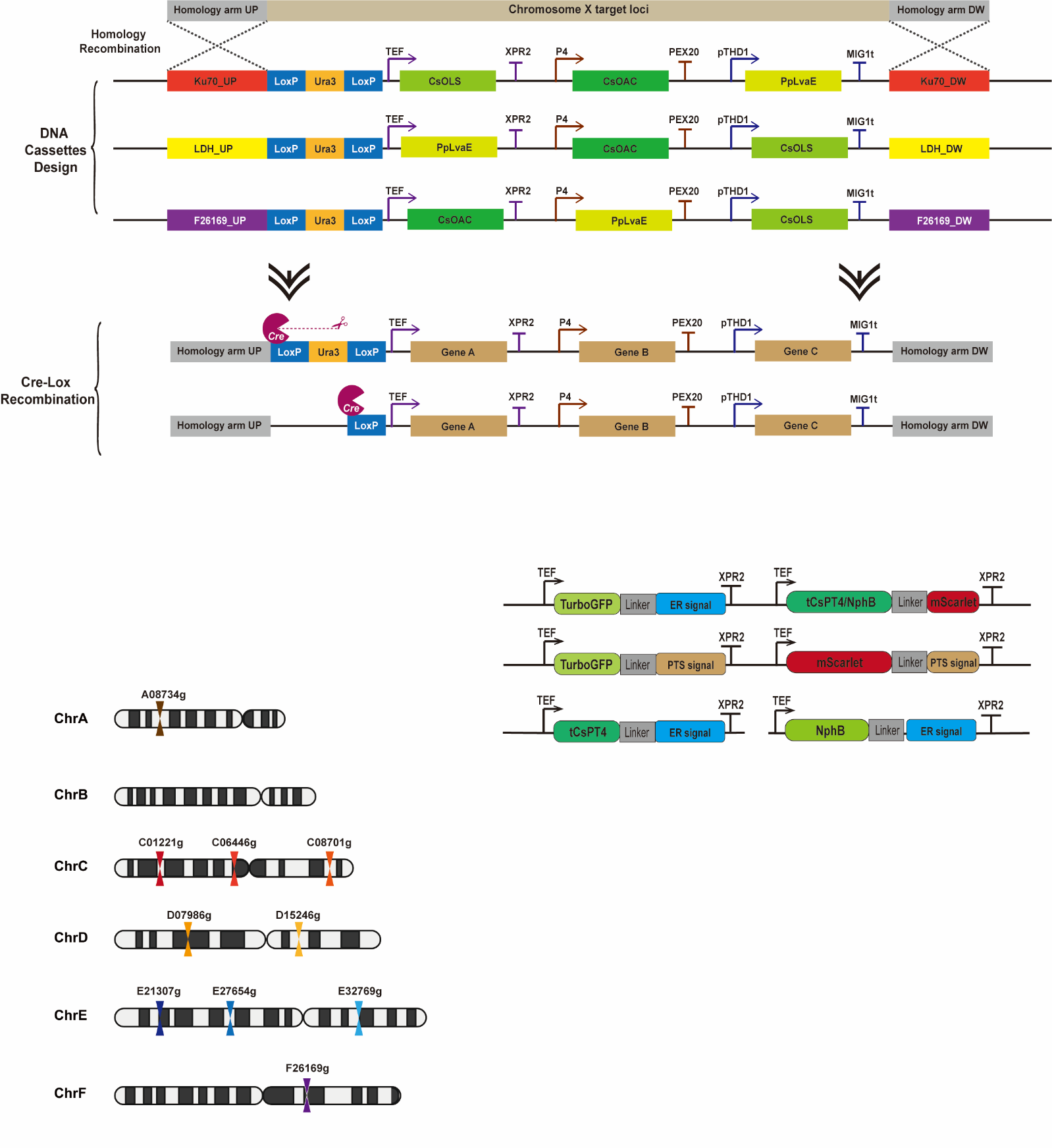

**Supplementary Figure S2 Yarrowia lipolytica chromosome locus used for integration in this study.**

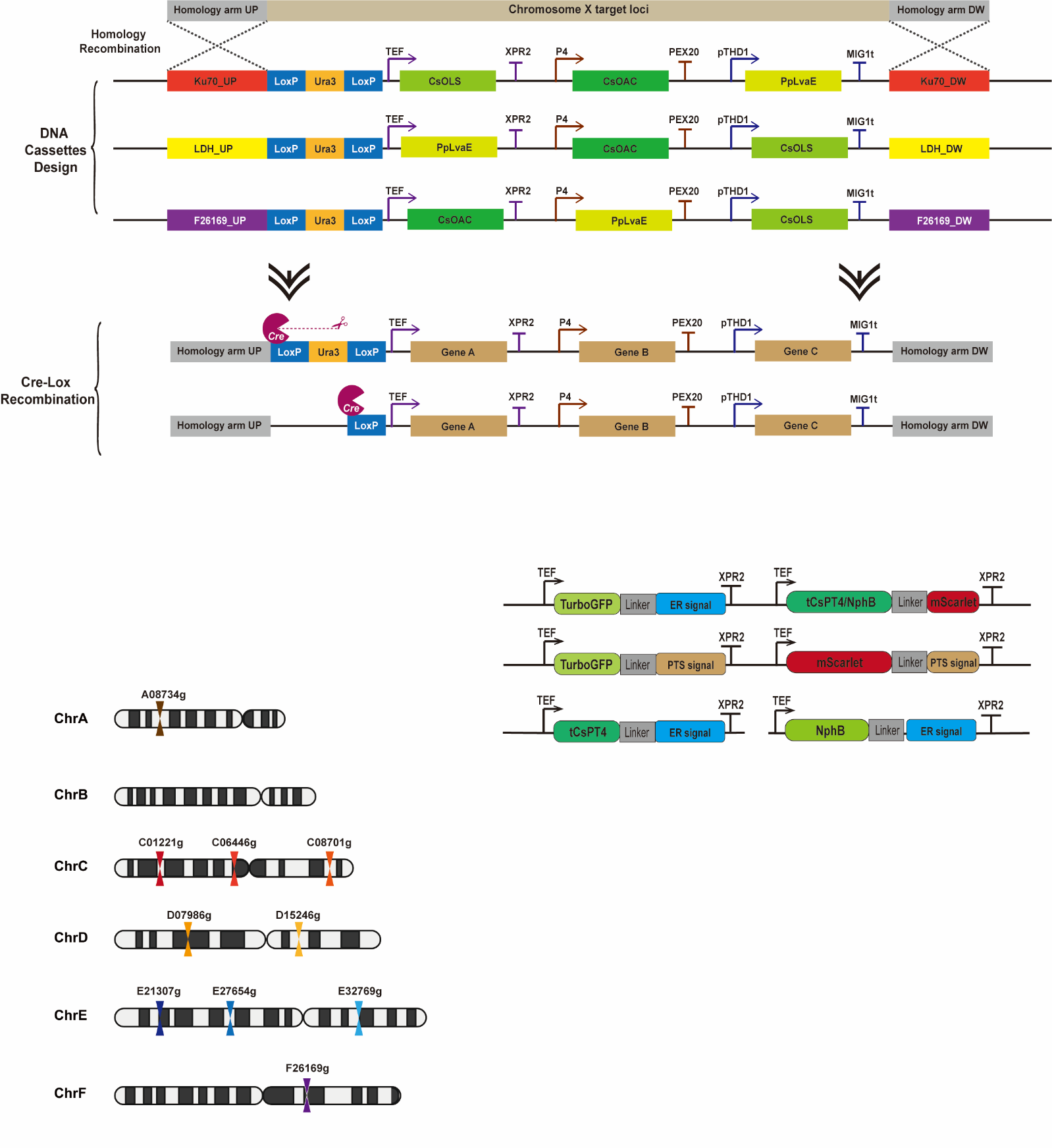

**Supplementary FigureS3. Olivetolic Acid LCMS Negative-ion mode MS2 fragments spectrum**

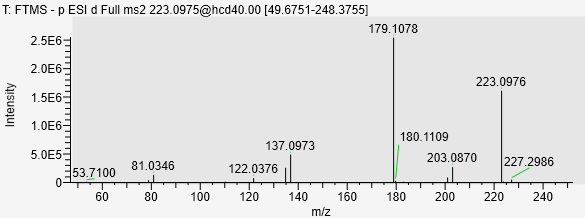

**Supplementary Figure S4 Schematic diagram of ER1/2, PTS signal, TurboGFP, and mScarlet fused construction in this study**

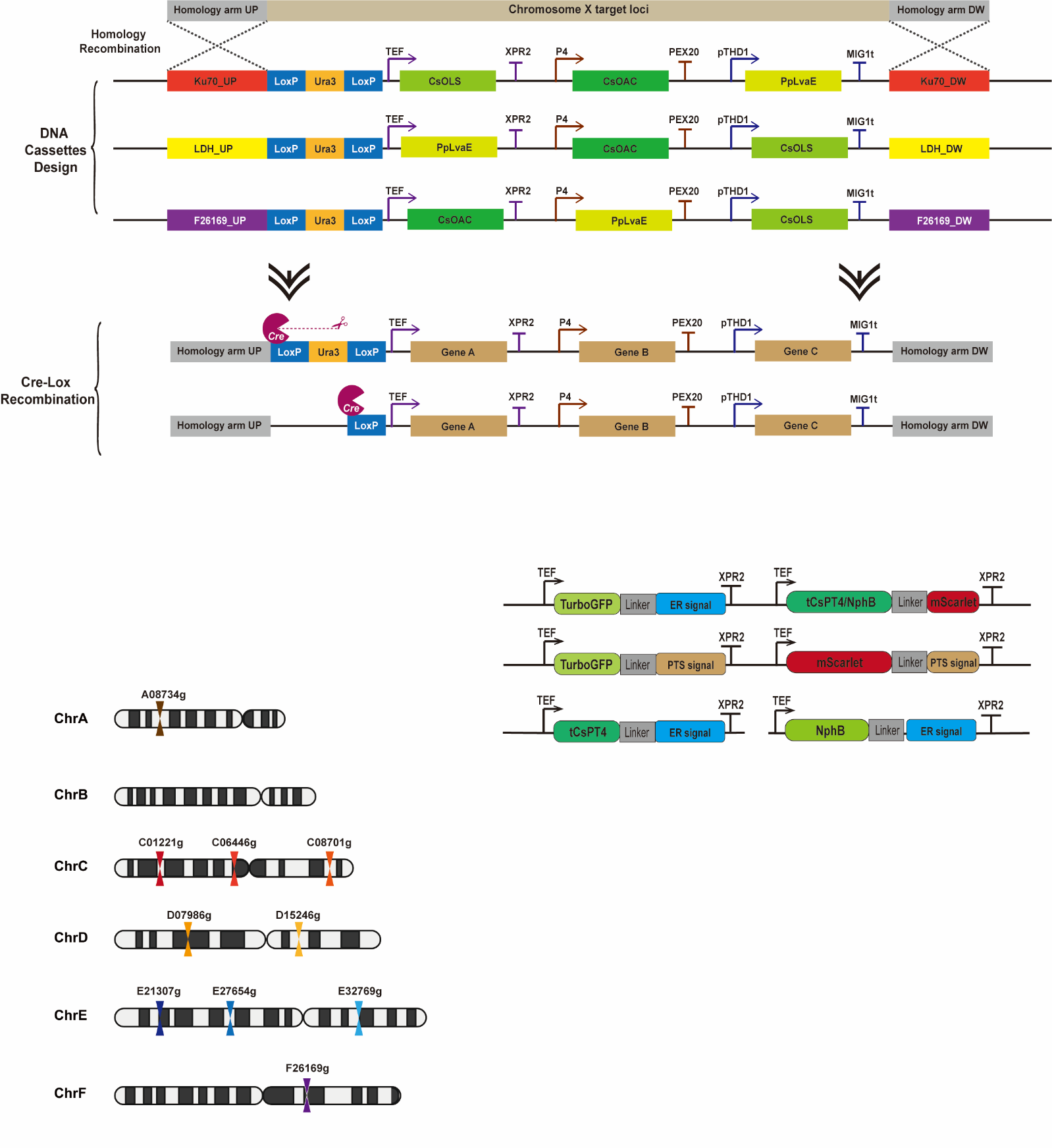

**Supplementary FigureS5. CBGA LCMS Negative-ion mode MS2 fragments spectrum**

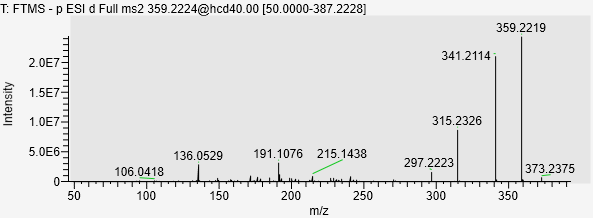

**Supplementary FigureS6. OSA LCMS Negative-ion mode MS2 fragments spectrum**

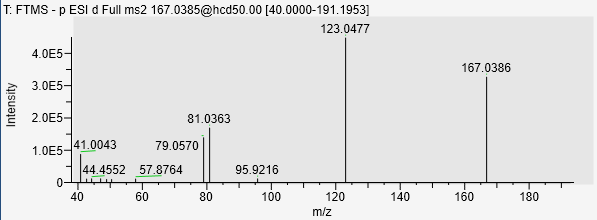

**Supplementary FigureS7. CBGOA LCMS Negative-ion mode MS2 fragments spectrum**

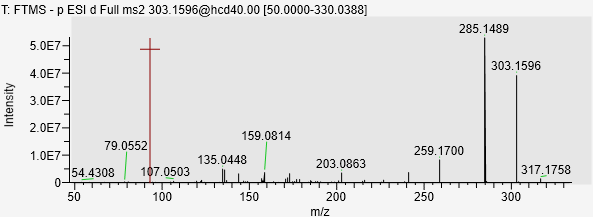

**Supplementary Table S1 The main engineered *Y. lipolytica* strains constructed in this study**

| **Strains** | **Relevant characteristics** | **Reference** |
| --- | --- | --- |
| po1g | W29ΔmatAΔxpr2-332Δaxp-2Δleu2-270 pBR platform | (Madzak et al., 2000) |
| po1fk | MatA, Leu-, Ura-, ΔAEP, ΔAXP, Suc+, ΔKu70 | (Gu et al., 2020a) |
| YX101 | Overexpressing CsOLS-CsOAC-PpLvaE in the genomic YALI0C08701g (Ku70) loci of po1fk | This study |
| YX102 | Overexpressing PpLvaE-CsOAC-CsOLS in the genomic YALI0E03212g (LDH) loci of YX101 | This study |
| YX103 | Overexpressing CsOAC-PpLvaE-CsOLS in the genomic YALI0F26169g loci of YX102 | This study |
| YX104 | Overexpressing ylACC1 in the genomic YALI0E32769g (DGA1) loci of YX103 | This study |
| YX105 | Overexpressing CsAAE1-tCsPT4 in the genomic YALI0C01221g loci of YX104 | This study |
| YX106 | Overexpressing CsAAE1-tCsPT4-PTS in the genomic YALI0C01221g loci of YX104 | This study |
| YX107 | Overexpressing EfmvaE-EfmvaS-ylERG20* in the genomic YALI0A08734g loci of YX105 | This study |
| YX108 | Overexpressing tCsPT4 in the 26s rDNA of YX107 | This study |
| YX109 | Overexpressing NphB* in the genomic YALI0E20977g(ARO8) loci of YX108 | This study |
| YX110 | Overexpressing NphB*-ER in the genomic YALI0E20977g(ARO8) loci of YX108 | This study |
| YX111 | Overexpressing tCsPT4-ER in the genomic YALI0E20977g(ARO8) loci of YX108 | This study |
| YX112 | Overexpressing ScERG20* in the genomic YALI0D00789g(GGPPS) loci of YX109 | This study |
| YX201 | Overexpressing ArmB in the genomic YALI0C08701g (Ku70) loci of po1fk | This study |
| YX202 | Overexpressing ylACC1 in the genomic YALI0E32769g (DGA1) loci of YX201 | This study |
| YX203 | Overexpressing ylACC1 in the genomic YALI0D07986g(DGA2) loci of YX201 | This study |
| YX204 | Overexpressing EfmvaE-EfmvaS-ylERG20* in the genomic YALI0A08734g loci of YX202 | This study |
| YX205 | Overexpressing tCsPT4 in the genomic YALI0C01221g loci of YX204 | This study |
| YX206 | Overexpressing NphB* in the genomic YALI0E20977g(ARO8) loci of YX205 | This study |
| YX207 | Overexpressing NphB*-ER in the genomic YALI0E20977g(ARO8) loci of YX205 | This study |
| YX208 | Overexpressing tCsPT4-ER in the genomic YALI0E20977g(ARO8) loci of YX205 | This study |

*** means mutation**

**Supplementary Table S2. Primers for sequencing and colony PCR used in this study**

| Name | Sequence |
| --- | --- |
| 26srDNA-fw | GCAGTCAAGCGTTCATAGCGACATTGC |
| 26srDNA-rvs | GGAGACCTGCTGCGGTTATCAGTACGAC |
| A0827-CKfw | ATTCGACTGAGCACGCCATCAGGTAGAC |
| A0827-CKRvs | TGATGGTATTGCACATTGACTCTTGATC |
| ADH-CK-Fw | CTCTGTGGTCCTGGTCCTTTGATTGTTC |
| ADH-CK-rvs | CAGGTGTACTGTAGCCACCCTGACATTA |
| ArmB_CK_FW | TCTATCAAGGTGATCCACCTGCCCG |
| ArmB_CK_Rvs | AGAGACTTGGGGTCCTTGAAGTGGT |
| armB-1-rvs | AGAAGACTTGGGGATGTACTCCTCCAGG |
| armB-2-fw | CCTGGAGGAGTACATCCCCAAGTCTTCT |
| armB-2-rvs | CCACCTCGGTGATCTTGGGCTTGGGAG |
| armB-3-fw | CTCCCAAGCCCAAGATCACCGAGGTGG |
| C0122-CKFw | ACGGTGTTTAAGCGGTCACTAACGG |
| C0122-CKrvs | CCGTCAAAATACCGGTCGAAATACAGTACA |
| CsAAE1-Rvs | GTGGCAGCTCCATAATTACACACGA |
| CsOLS-Rvs | AAAAATGAGATGCGTGATTTTCGACTTGG |
| D15246-CKFw | TGGTAGTTGACAGTGGCAATCTCAATGA |
| D15246-CKRvs | CTGCTGGTCTTCCACATGACTCAACTCT |
| DGA1_Fwd | ATCCGACCAGCACTTTTTGCAGTACTAACCGCAGACTATCGACTCACAATACTACAAGTCGCGAG |
| DGA1_Rvs | GTGGGGACAGGCCATGGAACTAGTCGGTACCTTACTCAATCATTCGGAACTCTGGGGCT |
| DGA1-GFP_Fwd | AGCCCCAGAGTTCCGAATGATTGAGGGCGGAGGCGGCGGAGGCGGAGGCGGAGGC |
| DLD2-DwCK-Rvs | GCTTGGTCTCCAATGCGATGGCGTCAAC |
| DLD2-UpCK-Fw | CGCCGAGATCAAGATTGCTTCTCGATCC |
| EfmvaE-fw | ATGAACCAAGACCGAGCCCTGGCCA |
| EfmvaE-Rvs | CACGGCAGACACTTGAGACAGAGAG |
| EfmvaS-fw | CTATCTCGGCAATCAACAACACGGT |
| EfmvaS-Rvs | GGATCGACATTCCGAGCTTCAGCCA |
| ER1-Oligo-Fw | TAGAGCAACCTCTGAAATTTGTGCTTACTGCGGCCGTCGTGCTCTTGACGACGTCGGTTCTTTGTTGTGTAGTATTTACA |
| ER1-Oligo-Rvs | GCACAAATTTCAGAGGTTGCTCTAATATGCCTCCGCCTCCGCCTCCGCCGCCTCCGCCTTCTTCACCGGCATCTGCATCCG |
| ER1-rvs | CGACGTCGTCAAGAGCACGA |
| ER2-Oligo-Fw | ACCAAAGTAAAGGTAGTGGTACATTGGTTGTCATATTGGCCATTTTAATGCTAGGTGTTGCTTATTATTTGTTGAACGAA |
| ER2-Oligo-Rvs | TGTACCACTACCTTTACTTTGGTTTTCAGAGGTAGAGCCTCCGCCTCCGCCTCCGCCGCCTCCGCCTTCTTCACCGGCATCTGCATCC |
| ERG20_F88W_Fwd | GCAGGCGTTTTGGCTCGTGTCGGAC |
| ERG20_F88W_Rvs | GTCCGACACGAGCCAAAACGCCTGC |
| ERG20_N119W_Fwd | CATGATTGCCATCTGGGATGCTTTCATG |
| ERG20_N119W_Rvs | CATGAAAGCATCCCAGATGGCAATCATG |
| F2616_DwChkR | GGCACCAGATTATCTTGATTGTGCG |
| F2616_UpChkF | GGGAGTAATAGAGGTGGAGTGAATGTTG |
| GGGGS-ER1-Fw | GGAGGTGGTGGATCTATATTAGAGCAACCTCTGAAATTTGTGCTTACTGC |
| GGGGS-NphB-Rvs | CTAATATAGATCCACCACCTCCGTCCTCGAGAGAATCAAATGCCTTGAG |
| GGGGS-tPT4-USA-Rvs | CTAATATAGATCCACCACCTCCGATAAAGACATAAACGAAGTATTCGGCG |
| Ku70_CKfwd | ATGGAATGGATTTCACATCTGGAGA |
| Ku70_CKrvs | TCACTTCCCATAGTACTTTTTGACC |
| LDH-DwCKR | CAAGAAGCACACCACATACTGTCATGGA |
| LDH-UpCKF | GAGAACCGAGTGTTCCTCAAGAAGTACA |
| M-CsOAC-rvs | CGCTCTCGAATGTAACCTCGACAATGT |
| M-CsOLS-fw | CGGTTGAACGAGTCGTTGTTAGAAGCG |
| M-CsOLS-rvs | TGCAGTACCGATAGCCAGGACGGATGC |
| mSca-Rvs | TGGGGACAGGCCATGGAACTAGTCGGTACCTTATTTATACAGCTCATCCATTCCTC |
| NphB-GT-CK-fw | CGAACGCTCGTCTATGGTCTGACAC |
| NphB-GT-CK-rvs | TAGCTGCATAGACACGCTCAACGTC |
| NphB-Line-Rvs | GTCCTCGAGAGAATCAAATGCCTTG |
| NphB-mSca-Fwd | CAAGGCATTTGATTCTCTCGAGGACGGCGGAGGCGGCGGAGGCGGAGGCGGAGGCGTTTCTAAGGGCGAAGCAGTTATTA |
| NphB-Mut1-fw | ACTCGGTGCAGTATATCACATCACGGACGTGCAG |
| NphB-Mut1-Rvs | GATATACTGCACCGAGTTTGTAGTATTCCTCCTTAGGGC |
| NphB-Mut2-fw | CTGCTTCAGTGTGATCAGCAACGACCCCAC |
| NphB-Mut2-Rvs | TGATCACACTGAAGCAGAGTCTATCAATTTTACCAGTC |
| POT1-CK-fw | TCATACCCTGGAGTGGAACCGCAAGATC |
| POT1-CK-rvs | TGACGTCATGATCGAGACGCTTATCGCC |
| POX4-DwChkR | AGCCGTTATATATATGCACCGAAGACC |
| POX4-UpChkF | TCAAATACTTTCCCCTTAGTAAACCTCGG |
| PT4-GFP_Fwd | CGAATATTTTGTCTACGTTTTCATCGGCGGAGGCGGCGGAGGCGGAGGCGGAGGC |
| PT4-Gly-GFP_Rvs | GGACAGGCCATGGAACTAGTCGGTACCCTATTCTTCACCGGCATCTGCATCCGGGGT |
| PT4-GT-pxs-rvs | GCTTAGAAGAGCCTCCGCCACCGATGAAAACGTAGACAAAATATTCGGCA |
| PT4-USA-Gly-fw | GCCGAATACTTCGTTTATGTCTTTATCGGCGGAGGCGGCGGAGGCGGAGGC |
| PT4-usa-pxs-fw | CGACCAGCACTTTTTGCAGTACTAACCGCAGGCAGGCTCCGATCAAA |
| PT4-usa-pxs-rvs | GCTTAGAAGAGCCTCCGCCACCGATAAAGACATAAACGAAGTATTCGGCG |
| pYL24P-Fw | CATTTATATAAGGGTCTGCATCGCCGGCT |
| pYL24P-Rvs | TACATCATCCCTCGGCCAAAGTTCATGT |
| pYL31P-Fw | ATAAACAGTGGCTCTCCCAATCG |
| pYL31P-Rvs | CCTTTTTCGTTGTTCCGACACCAACCTT |
| Scarlet-Rvs | AGAATGTCCCAGCTAAAGGGAAG |
| tCsPT4-GT-Fwd | CGACCAGCACTTTTTGCAGTACTAACCGCAGGCTGGTTCCGACCAGATCGAAGGA |
| tCsPT4-GT-Rvs | GCTTCCTTCGATCTGGTCGGAACCAGC |
| tCsPT4-mSca-Fwd | GCCGAATACTTCGTTTATGTCTTTATCGGCGGAGGCGGCGGAGGCGGAGGCGGAGGCGTTTCTAAGGGCGAAGCAGTTAT |
| TEF_Fwd | CAACTCACACCCGAAATCGT |
| TEF_Fwd_2 | AGAGACCGGGTTGGCGGCGCATTTG |
| TurboGFP-ER1-Fw | GGTTCTTTGTTGTGTAGTATTTACATGAGGTACCGACTAGTTC |
| TurboGFP-ER2-Fw3 | TGTTGCTTATTATTTGTTGAACGAATGAGGTACCGACTAGTTC |
| TurboGFP-ER-Fw | GGCTACTACAGCTCCGTGGT |
| TurboGFP-Line-Rvs | TTCTTCACCGGCATCTGCATCCGG |
| Urlp-Ura3-fw | CTAGGATAACTTCGTATAGCATACATTATACG |
| Urlp-Ura3-rvs | GTCGACTTGCATGCGCGGCCGC |
| XPR2_Rvs | GACAGGTAGTAGGAGGCAAG |
| XPR2_Rvs_2 | GGACACGGGCATCTCACTTGCATAT |
| XPR2_Urlp_FwD | ATGCGGCGTCTTTGTTCAT |
| ylACC_Down_Fwd | TGGACGAGGTTGTGGTTGACAAGAG |
| ylACC_Down_Rvs | GTGGGGACAGGCCATGGAACTAGTCGGTACCTCACAACCCCTTGAGCAGCTCAG |
| ylACC_UP_Rvs | CTCTTGTCAACCACAACCTCGT |
| ylACC2_Down_Fwd | CTGGAACCGGAGTGGACGAGGTTGTGGTTGACAAGAG |
| YliDGA1_ChkF | GTTTATGCATTCTGTTGGACCTTAGTCTG |
| YliDGA1_ChkR | GTTATCTACCACGATTTTTTGGTTTCTGAGGC |
| ylURA3-fw | GAGAACTCCCAGTACAAGGAGTTCC |
| ylURA3-rvs | TAGGTGAAGTCGTCAATGATGTCGATAT |

**Supplementary Sequence 1. The sequence of PpLvaE used in this study**

ATGGTTCCCACCCTCGAGCACGAACTCGCACCCAATGAGGCTAATCACGTCCCCCTGAGCCCCCTCTCTTTCCTGAAACGTGCCGCTCAAGTTTATCCTCAGAGAGATGCCGTTATTTACGGAGCTAGACGGTATTCCTATCGTCAACTCCACGAGCGATCGCGAGCACTCGCATCGGCACTCGAACGGGTGGGCGTGCAGCCCGGCGAAAGAGTCGCTATCCTGGCTCCCAATATTCCCGAAATGCTCGAAGCACACTATGGCGTCCCTGGCGCCGGCGCCGTGCTTGTTTGTATCAACATCCGTCTTGAGGGCCGAAGCATTGCCTTTATTCTTCGACATTGTGCTGCTAAAGTTCTCATTTGCGATAGAGAATTCGGAGCCGTGGCTAACCAGGCCCTCGCCATGCTGGACGCTCCTCCTCTCCTTGTGGGCATCGACGATGACCAAGCCGAGAGAGCAGACCTTGCACACGACCTGGACTATGAGGCATTCCTCGCTCAAGGTGATCCTGCCCGACCCCTGTCTGCCCCCCAGAACGAGTGGCAGTCGATCGCCATTAACTACACCTCCGGTACGACAGGCGATCCCAAGGGCGTTGTTCTCCACCACAGAGGCGCCTACCTTAATGCATGCGCCGGCGCACTGATCTTTCAGCTGGGTCCCAGATCCGTTTATCTGTGGACGCTGCCCATGTTCCATTGCAATGGATGGTCCCACACGTGGGCTGTCACACTTTCGGGAGGAACCCACGTCTGTCTGCGAAAAGTCCAGCCCGACGCAATTAACGCTGCAATTGCCGAGCATGCCGTGACACATCTTTCCGCAGCTCCTGTGGTCATGTCTATGCTTATTCATGCTGAGCACGCATCGGCTCCTCCTGTGCCCGTGTCCGTGATCACAGGCGGCGCAGCACCTCCTAGCGCCGTCATCGCCGCTATGGAGGCTCGAGGTTTTAACATCACCCACGCATACGGAATGACCGAGAGCTACGGTCCTTCGACACTGTGCCTGTGGCAGCCCGGTGTTGACGAACTCCCTCTGGAAGCCCGAGCCCAGTTCATGTCTCGACAGGGCGTTGCACATCCTCTGCTTGAAGAAGCCACAGTTCTGGACACTGATACCGGACGACCTGTCCCTGCAGATGGTCTGACGCTCGGAGAGCTTGTCGTCCGTGGAAACACCGTGATGAAAGGCTATCTTCATAATCCTGAGGCAACACGTGCCGCCCTCGCCAACGGTTGGCTGCACACAGGCGACCTTGCCGTGCTCCATCTGGACGGCTATGTGGAAATCAAAGACCGTGCAAAGGATATTATTATTTCCGGCGGTGAAAACATTTCTAGCCTTGAAATCGAAGAAGTTCTTTACCAACATCCTGAAGTGGTCGAAGCCGCCGTGGTGGCTAGACCCGACTCGAGATGGGGAGAAACGCCTCATGCTTTCGTCACACTTCGGGCAGATGCTCTGGCTTCCGGCGACGACCTGGTTCGGTGGTGTCGGGAGCGTCTTGCTCACTTTAAGGCCCCCCGTCATGTCTCCCTGGTGGATCTCCCTAAAACTGCTACCGGCAAAATCCAGAAGTTCGTTCTTCGAGAATGGGCCAGACAGCAGGAGGCTCAAATTGCAGATGCCGAACATTAA

**Supplementary Sequence 2. The sequence of CsOAC used in this study**

GCTGTTAAACACCTTATTGTTCTCAAATTCAAAGATGAGATTACCGAGGCTCAAAAAGAAGAATTTTTCAAGACGTATGTGAATCTGGTCAACATCATTCCCGCCATGAAAGACGTTTATTGGGGAAAAGATGTTACACAAAAGAATAAGGAAGAGGGCTATACTCACATTGTTGAGGTCACCTTCGAGTCGGTTGAGACTATTCAGGACTACATTATCCATCCCGCCCATGTTGGTTTCGGCGATGTTTATCGGAGCTTCTGGGAGAAACTGCTGATTTTTGATTATACTCCCCGGAAGTAA

**Supplementary Sequence 3. The sequence of CsOLS used in this study**

AATCATCTTCGGGCCGAGGGACCTGCCAGCGTGCTTGCCATTGGCACTGCTAACCCCGAGAACATCCTCCTTCAGGATGAATTCCCCGACTATTACTTCCGAGTCACCAAGTCCGAGCACATGACGCAGCTGAAAGAAAAATTTCGAAAGATTTGCGATAAATCCATGATTCGAAAGCGGAATTGTTTCCTGAACGAGGAACACCTCAAGCAGAACCCCCGTCTGGTGGAGCACGAGATGCAAACCCTTGACGCTAGACAAGACATGCTCGTGGTCGAAGTCCCTAAACTTGGAAAAGACGCCTGTGCAAAAGCAATCAAAGAATGGGGTCAGCCCAAGTCGAAAATCACGCATCTCATTTTTACATCCGCCTCGACGACTGACATGCCTGGAGCTGACTATCACTGCGCCAAGCTCCTGGGTCTTTCCCCTTCTGTCAAGAGAGTGATGATGTACCAGCTGGGTTGCTACGGAGGTGGCACGGTTCTTAGAATCGCAAAGGACATCGCTGAGAATAACAAGGGTGCACGGGTGCTCGCTGTGTGCTGTGATATTATGGCCTGCCTGTTTCGTGGCCCTTCTGAGTCGGACCTTGAGCTTCTGGTCGGCCAGGCCATTTTTGGAGATGGAGCAGCCGCTGTTATCGTTGGCGCCGAGCCTGATGAGAGCGTCGGTGAGCGACCTATTTTTGAACTTGTCAGCACTGGTCAAACAATCCTGCCTAACTCTGAAGGAACGATTGGAGGCCATATCCGAGAGGCTGGACTGATCTTCGATCTGCATAAGGACGTTCCCATGCTTATTTCGAATAATATTGAAAAGTGTCTCATTGAGGCCTTTACTCCCATCGGCATCTCGGACTGGAATTCCATCTTTTGGATCACTCACCCCGGTGGTAAAGCAATCCTTGACAAAGTTGAGGAAAAACTGCACCTTAAAAGCGATAAGTTTGTTGACTCGCGTCACGTGCTCTCTGAGCATGGTAACATGTCGTCCTCCACTGTTCTGTTCGTCATGGATGAGCTTCGGAAACGTTCGCTTGAGGAAGGAAAATCCACCACAGGAGATGGTTTCGAGTGGGGCGTGCTCTTTGGTTTCGGACCCGGCCTCACGGTTGAGCGGGTGGTCGTGCGATCGGTGCCCATTAAGTACTAG

**Supplementary Sequence 4. The sequence of CsAAE1 used in this study**

GGCAAAAATTATAAATCCCTTGACTCGGTGGTTGCTTCGGACTTTATCGCACTGGGTATCACCTCTGAGGTTGCCGAAACGCTCCACGGTAGACTCGCCGAAATCGTGTGTAATTATGGAGCTGCCACGCCCCAAACATGGATTAATATTGCCAACCACATCCTTAGCCCCGATCTCCCCTTCTCTCTTCACCAGATGCTGTTTTATGGCTGCTACAAGGATTTCGGACCCGCACCCCCCGCATGGATCCCTGACCCCGAAAAAGTTAAGTCTACCAACCTTGGTGCCCTCCTTGAGAAGAGAGGCAAGGAATTCCTGGGAGTTAAATATAAAGACCCTATCTCTTCCTTTAGCCATTTTCAGGAGTTCTCTGTTAGAAACCCCGAGGTCTACTGGCGGACAGTTCTCATGGATGAAATGAAGATCTCCTTTAGCAAAGACCCCGAGTGCATCCTCCGACGGGATGACATTAATAATCCTGGAGGTTCGGAATGGCTGCCTGGAGGCTATCTGAACTCTGCTAAGAACTGCCTGAACGTTAATTCCAACAAAAAGCTTAACGATACAATGATTGTCTGGCGTGACGAGGGTAATGACGATCTGCCCCTTAATAAGCTTACCCTCGACCAACTTCGGAAGCGAGTTTGGCTGGTTGGTTATGCTCTTGAGGAGATGGGCCTTGAAAAGGGCTGTGCTATCGCTATTGACATGCCCATGCACGTGGATGCAGTCGTTATCTATCTCGCAATTGTTCTCGCCGGTTACGTCGTCGTTTCGATTGCCGATTCGTTTAGCGCTCCTGAAATCTCTACACGACTGAGACTCAGCAAGGCCAAGGCAATCTTCACACAAGATCACATTATTCGAGGTAAGAAACGGATTCCCCTTTACAGCCGAGTGGTGGAGGCCAAGAGCCCTATGGCAATTGTGATCCCCTGCTCGGGATCGAATATTGGCGCTGAACTTCGGGATGGTGACATCTCTTGGGATTATTTCCTTGAACGGGCAAAAGAGTTTAAAAATTGTGAATTTACAGCACGAGAACAACCCGTTGACGCATATACAAACATTCTCTTCAGCTCTGGCACGACTGGAGAACCCAAAGCCATTCCTTGGACTCAGGCAACCCCCCTTAAGGCTGCAGCTGACGGTTGGTCTCATCTGGATATCAGAAAGGGCGATGTGATTGTTTGGCCTACAAATCTCGGATGGATGATGGGACCCTGGCTCGTCTATGCTTCGCTGCTTAATGGCGCTTCTATCGCCCTTTACAATGGTTCTCCTCTGGTCTCCGGATTTGCCAAATTCGTGCAAGATGCTAAGGTCACAATGCTTGGTGTGGTGCCTTCTATCGTTCGGTCGTGGAAAAGCACGAACTGCGTTTCTGGCTACGACTGGAGCACTATTCGTTGCTTCTCTTCCAGCGGCGAGGCCTCCAACGTTGATGAGTATCTGTGGCTGATGGGACGAGCTAATTACAAACCTGTCATCGAGATGTGTGGTGGCACAGAAATCGGCGGTGCTTTTTCTGCCGGTTCGTTCCTCCAGGCCCAGTCTCTTTCCAGCTTTAGCTCCCAATGTATGGGTTGTACGCTGTACATTCTCGATAAGAACGGTTACCCCATGCCTAAGAATAAGCCCGGTATCGGCGAGCTTGCACTGGGTCCTGTTATGTTTGGTGCCTCGAAAACGCTCCTGAACGGCAATCATCACGATGTTTACTTTAAAGGCATGCCTACCCTCAACGGAGAGGTTCTGAGACGACACGGAGATATCTTTGAGCTTACATCGAATGGATATTATCATGCACACGGAAGAGCAGATGACACGATGAATATTGGTGGAATTAAAATTAGCTCGATTGAAATTGAGCGAGTCTGTAACGAGGTCGATGACCGAGTCTTCGAAACAACAGCTATTGGCGTTCCTCCCCTTGGCGGCGGTCCTGAGCAACTGGTTATCTTCTTTGTCCTCAAAGATTCTAATGACACGACGATCGACCTTAACCAGCTGCGACTGTCTTTTAATCTGGGTCTGCAGAAAAAACTTAACCCTCTTTTCAAAGTCACTCGGGTCGTGCCTCTGTCGAGCCTTCCTCGGACAGCCACCAATAAAATCATGAGACGTGTCCTTCGGCAGCAATTTAGCCATTTTGAGTAA

**Supplementary Sequence 5. The sequence of EfmvaE used in this study**

AAGACCGTGGTGATCATCGACGCCCTGCGAACCCCCATCGGCAAGTACAAGGGCTCTCTGTCTCAAGTGTCTGCCGTGGACCTGGGCACCCACGTGACCACTCAGCTGCTGAAGCGACACTCTACCATCTCTGAAGAAATCGACCAAGTGATCTTCGGCAACGTGCTGCAAGCCGGCAACGGACAGAACCCCGCCCGACAGATCGCCATCAACTCTGGCCTGTCTCACGAGATCCCCGCCATGACTGTGAACGAGGTGTGTGGCTCTGGCATGAAGGCCGTGATCCTGGCCAAGCAGCTGATTCAGCTGGGCGAGGCCGAGGTGCTGATCGCCGGCGGCATCGAGAACATGTCTCAAGCCCCCAAGCTGCAGCGATTCAACTACGAGACCGAGTCTTACGACGCCCCCTTCTCTTCTATGATGTACGACGGCCTGACCGACGCCTTCTCTGGCCAAGCCATGGGCCTGACCGCCGAGAACGTGGCCGAGAAGTACCACGTGACCCGAGAGGAGCAAGATCAGTTCTCTGTGCACTCTCAGCTGAAGGCCGCCCAAGCCCAAGCCGAGGGCATCTTCGCCGACGAGATCGCCCCCCTGGAGGTGTCTGGCACCCTGGTGGAGAAGGACGAGGGCATCCGACCCAACTCTTCTGTGGAGAAGCTGGGCACCCTGAAGACCGTGTTCAAGGAGGACGGCACCGTGACCGCCGGCAACGCCTCTACCATCAACGACGGCGCCTCTGCCCTGATCATCGCCTCTCAAGAGTACGCCGAGGCCCACGGCCTGCCCTACCTGGCCATCATCCGAGACTCTGTGGAGGTGGGCATCGACCCCGCCTACATGGGCATCTCTCCCATCAAGGCCATTCAGAAGCTGCTGGCCCGAAATCAGCTGACCACCGAGGAGATCGACCTGTACGAGATCAACGAGGCCTTCGCCGCCACCTCTATCGTGGTGCAGCGAGAGCTGGCCCTGCCCGAGGAGAAGGTGAACATCTACGGCGGAGGCATCTCTCTGGGCCACGCCATCGGAGCTACCGGCGCCCGACTGCTGACCTCTCTGTCTTATCAGCTGAATCAGAAGGAGAAGAAGTACGGCGTGGCCTCTCTGTGTATCGGCGGAGGCCTGGGCCTGGCCATGCTGCTGGAGCGACCTCAGCAGAAGAAGAACTCTCGATTCTATCAGATGTCTCCTGAGGAGCGACTGGCCTCTCTGCTGAACGAGGGACAGATCTCTGCCGACACCAAGAAGGAGTTCGAGAACACCGCCCTGTCTTCTCAGATCGCCAACCACATGATCGAGAATCAGATCTCTGAGACCGAGGTGCCCATGGGCGTGGGCCTGCACCTGACCGTGGACGAGACCGACTACCTGGTGCCCATGGCCACCGAGGAGCCCTCTGTGATCGCCGCCCTGTCTAACGGCGCCAAGATCGCCCAAGGCTTCAAGACCGTGAATCAGCAGCGACTGATGCGAGGACAGATCGTGTTCTACGACGTGGCCGACGCCGAGTCTCTGATCGACGAGCTGCAAGTGCGAGAGACCGAGATCTTTCAGCAAGCCGAGCTGTCTTACCCCTCTATCGTGAAGCGAGGCGGCGGACTGCGAGACCTGCAGTACCGAGCCTTCGACGAGTCTTTCGTGTCTGTGGACTTCCTGGTGGACGTGAAGGACGCCATGGGCGCCAACATCGTGAACGCCATGCTGGAGGGCGTGGCCGAGCTGTTCCGAGAGTGGTTCGCCGAGCAGAAGATCCTGTTCTCTATCCTGTCTAACTACGCCACCGAGTCTGTGGTGACCATGAAGACCGCCATCCCTGTGTCTCGACTGTCTAAGGGCTCTAACGGCCGAGAGATCGCCGAGAAGATCGTGCTGGCCTCTCGATACGCCTCTCTGGACCCCTACCGAGCCGTGACCCACAACAAGGGCATCATGAACGGCATCGAGGCCGTGGTGCTGGCCACCGGCAACGACACCCGAGCCGTGTCTGCCTCTTGTCACGCCTTCGCCGTGAAGGAGGGCCGATACCAAGGCCTGACCTCTTGGACCCTGGACGGCGAGCAGCTGATCGGCGAAATCTCTGTGCCCCTGGCCCTGGCCACCGTGGGAGGCGCCACTAAAGTCCTGCCCAAGTCTCAAGCCGCTGCCGACCTGCTGGCCGTGACCGACGCCAAGGAACTGTCTCGAGTGGTGGCCGCCGTGGGCCTGGCTCAGAATCTGGCCGCCCTGCGAGCCCTGGTGTCTGAGGGCATTCAGAAGGGCCACATGGCCCTGCAAGCCCGATCTCTGGCCATGACCGTGGGAGCCACCGGCAAAGAGGTGGAGGCCGTGGCTCAGCAGCTGAAGAGACAGAAAACCATGAACCAAGACCGAGCCCTGGCCATCCTGAACGACCTGCGAAAGCAGTAA

**Supplementary Sequence 6. The sequence of EfmvaS used in this study**

ACGATCGGTATCGACAAAATCAGCTTCTTCGTGCCTCCCTATTACATTGATATGACAGCCCTGGCTGAAGCTCGGAATGTCGATCCTGGCAAATTTCATATCGGCATCGGACAGGACCAAATGGCCGTTAATCCTATTAGCCAAGATATTGTTACATTTGCAGCAAACGCTGCCGAAGCTATTCTGACCAAGGAAGATAAGGAAGCTATTGATATGGTTATTGTGGGAACAGAAAGCTCCATTGACGAGTCGAAGGCCGCAGCCGTCGTTCTTCATCGGCTTATGGGTATTCAGCCCTTTGCTCGATCTTTTGAGATTAAAGAAGCCTGTTACGGAGCCACTGCAGGTCTGCAACTCGCTAAGAATCATGTTGCCCTGCATCCCGATAAAAAGGTCCTGGTCGTTGCAGCAGACATTGCTAAGTATGGACTGAATTCCGGTGGTGAGCCCACTCAGGGCGCAGGTGCTGTCGCTATGCTTGTTGCAAGCGAGCCTCGAATTCTGGCACTGAAAGAAGATAATGTGATGCTTACACAGGACATCTACGACTTTTGGCGTCCCACAGGTCACCCTTACCCCATGGTTGATGGACCTCTGTCGAACGAAACATATATCCAATCTTTCGCTCAGGTGTGGGATGAACATAAAAAAAGAACAGGCCTTGATTTCGCTGATTATGATGCTCTGGCATTTCACATCCCTTATACGAAAATGGGCAAAAAGGCCCTGCTGGCTAAGATTTCTGACCAGACGGAGGCAGAGCAAGAACGAATCCTCGCACGTTACGAAGAATCGATCGTCTATTCTCGGCGTGTGGGCAATCTTTATACAGGCTCCCTCTACCTGGGCCTGATTTCCCTGCTGGAAAACGCAACAACACTCACCGCTGGCAATCAAATCGGCCTCTTCTCTTACGGTTCCGGCGCTGTCGCAGAATTTTTTACAGGTGAACTTGTTGCTGGATATCAGAACCACCTGCAAAAAGAAACCCACCTGGCACTTCTCGATAACCGTACAGAGCTGTCCATTGCTGAGTATGAAGCAATGTTCGCAGAGACGCTGGACACTGACATTGACCAGACACTGGAGGACGAACTCAAGTATTCTATCTCGGCAATCAACAACACGGTTAGAAGCTACCGGAACTAG

**Supplementary Sequence 7. The sequence of CsPT4 used in this study**

GGACTGAGCCTTGTTTGCACATTCTCTTTTCAGACAAATTATCATACTCTTCTCAACCCTCATAATAAAAATCCCAAGAATAGCCTTCTGTCCTATCAACACCCTAAGACGCCCATTATCAAGTCCTCTTACGATAATTTTCCCTCTAAATACTGCCTTACAAAGAATTTCCACCTCCTCGGACTCAATTCCCATAACCGAATCTCCTCGCAATCTCGTAGCATCCGGGCTGGTTCCGACCAGATCGAAGGAAGCCCTCATCATGAATCGGATAATAGCATCGCTACTAAGATCCTGAACTTTGGTCACACGTGCTGGAAACTTCAGCGTCCCTATGTCGTCAAGGGCATGATTTCCATCGCTTGCGGTCTCTTTGGACGGGAACTGTTTAACAACAGACATCTCTTTTCTTGGGGTCTTATGTGGAAGGCCTTCTTCGCACTTGTCCCCATCCTCTCCTTCAATTTTTTTGCAGCCATCATGAATCAAATCTATGACGTGGACATTGACCGTATCAACAAGCCTGACCTTCCTCTTGTGTCTGGAGAAATGTCGATTGAAACCGCATGGATCCTTTCGATCATCGTGGCCCTGACTGGCCTCATCGTCACGATTAAGCTGAAATCTGCTCCTCTCTTTGTGTTCATTTACATTTTTGGCATCTTCGCTGGTTTCGCCTATTCCGTCCCCCCCATTCGATGGAAACAATATCCTTTCACAAACTTCCTTATCACCATTTCCTCCCACGTTGGCCTCGCTTTCACGTCCTATTCTGCTACAACTAGCGCACTTGGCCTGCCCTTTGTGTGGCGTCCCGCTTTCTCGTTTATTATCGCTTTTATGACAGTCATGGGCATGACCATTGCATTTGCAAAGGACATTTCCGACATTGAAGGCGACGCTAAGTACGGAGTTAGCACGGTGGCTACGAAACTTGGAGCCCGGAATATGACTTTTGTGGTTTCCGGAGTTCTGCTCCTGAACTATCTGGTGTCGATCTCTATCGGAATCATCTGGCCTCAAGTTTTCAAGTCCAACATCATGATCCTCTCGCACGCTATCCTGGCCTTCTGTCTTATCTTTCAAACTAGAGAGCTTGCTCTTGCCAATTACGCAAGCGCACCTTCCCGGCAATTTTTTGAGTTCATTTGGCTTCTGTACTATGCCGAATATTTTGTCTACGTTTTCATCTAG

**Supplementary Sequence 8. The sequence of NphB used in this study**

TCGGAAGCCGCCGACGTTGAGCGTGTCTATGCAGCTATGGAGGAGGCCGCAGGACTCCTCGGTGTCGCATGTGCCAGAGACAAAATCTATCCCCTCCTTTCTACCTTCCAGGATACTCTGGTGGAAGGAGGTTCTGTGGTTGTGTTTTCCATGGCCAGCGGCAGACATAGCACCGAACTTGATTTCTCGATTAGCGTCCCTACCAGCCACGGCGACCCTTACGCAACGGTCGTGGAGAAAGGACTTTTTCCCGCAACCGGACATCCCGTGGACGATCTTCTCGCAGACACCCAGAAACACCTTCCTGTGAGCATGTTCGCTATCGACGGTGAAGTGACAGGCGGATTTAAGAAGACCTATGCTTTCTTTCCCACGGACAATATGCCTGGTGTGGCAGAACTGTCGGCCATCCCCTCGATGCCTCCCGCTGTGGCCGAAAACGCTGAACTTTTCGCACGATATGGTCTTGATAAGGTCCAGATGACAAGCATGGATTATAAAAAGCGGCAAGTCAACCTTTATTTCTCTGAACTGTCTGCCCAGACCCTCGAAGCTGAATCGGTGCTTGCACTGGTGCGTGAGCTCGGACTCCATGTTCCCAATGAACTTGGACTCAAATTCTGCAAAAGATCTTTTAGCGTTTACCCTACCCTGAACTGGGAGACTGGTAAAATTGATAGACTCTGCTTCGCAGTGATCAGCAACGACCCCACACTTGTGCCCTCCAGCGACGAGGGAGATATTGAAAAATTTCACAATTATGCTACTAAGGCCCCCTATGCTTATGTCGGTGAGAAACGAACGCTCGTCTATGGTCTGACACTTAGCCCTAAGGAGGAATACTACAAACTCTCCGCAGCCTATCACATCACGGACGTGCAGCGAGGTCTTCTCAAGGCATTTGATTCTCTCGAGGACTAA

**Supplementary Sequence 9. The sequence of ScERG20 used in this study**

GCTTCAGAAAAAGAAATTAGGAGAGAGAGATTCTTGAACGTTTTCCCTAAATTAGTAGAGGAATTGAACGCATCGCTTTTGGCTTACGGTATGCCTAAGGAAGCATGTGACTGGTATGCCCACTCATTGAACTACAACACTCCAGGCGGTAAGCTAAATAGAGGTTTGTCCGTTGTGGACACGTATGCTATTCTCTCCAACAAGACCGTTGAACAATTGGGGCAAGAAGAATACGAAAAGGTTGCCATTCTAGGTTGGTGCATTGAGTTGTTGCAGGCTTACTGGTTGGTCGCCGATGATATGATGGACAAGTCCATTACCAGAAGAGGCCAACCATGTTGGTACAAGGTTCCTGAAGTTGGGGAAATTGCCATCTGGGACGCATTCATGTTAGAGGCTGCTATCTACAAGCTTTTGAAATCTCACTTCAGAAACGAAAAATACTACATAGATATCACCGAATTGTTCCATGAGGTCACCTTCCAAACCGAATTGGGCCAATTGATGGACTTAATCACTGCACCTGAAGACAAAGTCGACTTGAGTAAGTTCTCCCTAAAGAAGCACTCCTTCATAGTTACTTTCAAGACTGCTTACTATTCTTTCTACTTGCCTGTCGCATTGGCCATGTACGTTGCCGGTATCACGGATGAAAAGGATTTGAAACAAGCCAGAGATGTCTTGATTCCATTGGGTGAATACTTCCAAATTCAAGATGACTACTTAGACTGCTTCGGTACCCCAGAACAGATCGGTAAGATCGGTACAGATATCCAAGATAACAAATGTTCTTGGGTAATCAACAAGGCATTGGAACTTGCTTCCGCAGAACAAAGAAAGACTTTAGACGAAAATTACGGTAAGAAGGACTCAGTCGCAGAAGCCAAATGCAAAAAGATTTTCAATGACTTGAAAATTGAACAGCTATACCACGAATATGAAGAGTCTATTGCCAAGGATTTGAAGGCCAAAATTTCTCAGGTCGATGAGTCTCGTGGCTTCAAAGCTGATGTCTTAACTGCGTTCTTGAACAAAGTTTACAAGAGAAGCAAATAG

**Supplementary Sequence 10. The sequence of ArmB used in this study**

TCTCCCTCTCTGGTGGTGCCCGTGTTCGCCGGCCACGGCACTACCGCCATCAACTCTACCTCTCTGCGAGAGCGAGCTGTGACCGACGCCTCTTCTTCCTCTGGCGCCCTGCTCCTGGACGCCTGTTACTACGCCTTCAACGTGGAGCTGTCTACCCTGTCTCCCTCTGAGGCCTCTGCCGTGGGCATCAACCCCGACCACTTCAAGGACCCCAAGTCTCTGCTCCTGCTGCCCTCTCACGAGCACTACTTCACCAACTCTGTGGTGACCGCCGCCACCCTGTTCCTGGTGCAGACCCTGCGATACCTGGCCTCTGTGCAAGCCTCTTCCTCTGCCTCTTTCGCTTCCACCCTGCAGATGAACTCTGAGCACGGCCTGGGCATCGTGGGCTTCTCTTCTGGCATCCTGCCCGCCTGTGTGGTGGGCTCTTCTGAGACCACCCTGGAGTTCATCTCTAACGCCGTGGAGACCTTCCGACTGGCCTTCTGGATCGGCGTGCGACTGCAAGTGCACAAGGCCTCTGTGGAGACCCCCGAGCTGCTGGGCGAGTCTCCCCTGCCCTGGTCTCTGGCCTTTCTGTCTATGTCTCCCGCCGCTGCCGAGTCTGCCATTCAGTCTTTCCACAAGTCTTTCGAGGGCACCCCTGAGCTGCGAGTGACCTCTGTGGTGTCTGAGACCTCTGTGACCATCTCTGGCCGACCCGACATTCTGGCCGCCTTTGCTGCTCAGCTGCCCCCCTCCGGCCCCGTGCACAAGACCACCGTGGACGCCCTGTATCACTCTTCCTCTCACCACGACGGCGTGCGATCTCAAGTGCTGGCCGATGTGATCCGACGAAACATCCGATTCCCCACCCACGCCGACATCAAGATTCCCGTGCGATCCACCTACTCTGGCGAGCTCCTCAACAAGTCTCCCGAGGGCTCTGCTTCTTTCGTCGAGCAAGTGATCGACATGATCCTGACTCAGCCCGTGAACTGGGACAAGGTGACCGAGGCCCTCGTGAGAGCCGCCCCTGAGGCTGAGGTGGTGCACCTGCTGAACTTCGGCCCCGGCGCCGGCCTGACCAAGGGCATCGAGCGATACTTCCCCTCTGGCAAGGTGTCCTCTATTGACCTGTCTACCGAGGCCGTGCACACTTCTACCCTCCAAATGCCCTCTTCTGTGCAAGAGCCCATCGCCATCTGTGGCATGTCTGTGAACATGCCCGGCGCTCAGTCTGTGGCTAAACTGTGGGAGGTGCTGGAGAAGGGCATCAACACCGTGTCTGAGGTGCCCGAGCACCGATTCAAGGTGTCTGACTACAACGACCCCAAGAAAAAGTCTCGAACCATGGCCGCCCACACCGGCAACTTCATCGACGAGCCCGATGCTTTCGACAACAAGTTCTTCAACATCTCTCCCCGAGAGGCCCGATCTATGGACCCTCAGCAGCGAGTCCTGCTGCATACTGCCTACGAGGCTCTGGAGGACGCCGGCTACGTGCCCAACTCTACCCCCACCAACAATCCCGAGACCTTCGGCTGTTACGTGGGCGTGGCCACCAACGACTACGTGCAGAACCTGCGAAACGACATCGACGTGTACTACTCTACCGGCACCCTGCGAGCCTTCCTGTCTGGCCGAATCTCTTACGCCCTGCAGTTCTCTGGCCCCTCTATCGTGGTGGACACCGCCTGTTCTTCTTCCCTGATCGCCGTGTACCAAGCCTGTCGAGCCCTGATGAACCGAGACTGTAACGCCGCCGTGGCCGGCGGCGTGAACGTGATCGGCTCTCCCGACATGTTCCTGGGACTGGACCGAGGCCACTTCCTGTCTCCCACCGGACAGTGTAAGGCCTTCGACGCCTCTGCCGACGGCTACTCTCGATCTGAGGGCTGTGGCATCTTCGTGCTGAAGCGACTGTCTGATGCCGTGGCCGAGAACGATCAGATCCTGGGCGTGATCCGAGGCGTGGAGGTGAATCAGTCTGGCAACGCCTACTCTATCACCCGACCCCACGCCCCCACCCAAGAGAACCTGTTCACTCAGACCCTGGAGCGATCTGGCCTGGACGCCTCTCGAATCTCTGTGGTGGAGGCCCACGGCACTGGCACCCAAGCCGGCGACCCCATCGAGCTGGAGTCTATCCGAGGCATCTTCGCCAAGAACCGAAAGACCAATAACCCCCTGCACATCACCTCTGTGAAGGCCAATATCGGCCACCTGGAGGCCGCTTCTGGCGCCGCTGCCCTGGCTAAACTGCTCCTGATGCTGCGACACCGAACCATCCCCCGACTGATCTCTCTGAAGAACCTGAACCCCCGAATCAAGCCCCTGGCCTCTGACAACGTGATCATCGACACCAAGCAAGTGGCCTGGGCCGTGCCCGACGAGTCTCTGCCCCGAGTGGCCCTGCTGAACAACTTCGGCGCCGCCGGCTCTAACGGCGCCCTCCTCCTGGAGGAGTACATCCCCAAGTCTTCTGAGAAGATTGAGGTGTCTTCTACCTTCATCGTGGGCCTGTCTGCCAAGAACGAGCAAGCCCTGGTGGACCTCCGAGCCTCTTACATCGAGTACCTGCGATCTCCCGCCTCTGCCGGCGTGTCTCTGGCCGACATCGCCTACACCGCCACCGCCCGAAGACGAATCTTCTCCCACCGATTCGCCGTGACCGTGAAGTCTAAGGAGGAGCTGGCCCACAAGCTGGAGCTGGCCTCTGGCAAGACCGTGTCTGACAAGGCCCCCGGCAAGGTGGTGTTCGTGTTCTCTGGCCAAGGCGGACAGTACCTGGGCATGGGCTCTGCCCTGTACAAGACCTCTACCCTCTTCAAGTCTGCCATCGACGAGTGTGAGTACTTCCTGAAAAAGAACAACTTCCCCGGCGTGCTGCCCATCATCACCTCTGACGGCGAGTCTTCTGAGCTGACCCCCGTCGAGGAATTCGAAGCCAACCAAGCCGCCATTTTCGCCCTCGAGTACGGCCTCGCCAAGCTGTGGATGTCTTGGGGCGTGACCCCCACCGCCGTGGTGGGCCATTCTCTGGGAGAGTACGCCGCCCACGTGGTGGCCGGCGTGCTGTCTCTGGAGTCTGCCCTGACCCTGGTGGCCCACCGAGTGCGAATCATGATGCGAACCTGTGAGCTGGACACCACCGGCATGATCGCCATCAACCTGGGCTCTGGCGCCGTCACCGACATTCTCTCTTCCTCTCCCGACTTCTCTGGCATCTCTATCGCCTGTTACAACTCTGCCACCGACTGTGTGGCCTCTGGCGCCATCGGACAGCTGGACGCCCTGAAAGCCCACCTGGACAAGAACGTGCACTGTAAGTCTGTGCGACTGAAGGTGCCCTTCGGCTACCACTCTTCCGCCATGCAGCCCCTGCTCGAGGAGTTCGGCGCCCTGGCCAAGCGAGTGACCGTGCACGCCCCCAAGATCCCCGTGATCTCCAACCCCCTGGGCCGAGTGGTGCGAGAGGGCGACAAGTCTGCCTTCAACGCCGAGTACTACCTGTCTCACTGTGCCGACCCCGTGCAGTTCGAGTCCGGCATCTCTGCCCTCATCGACGACGTGTCTTTTATGGACATTGCCGCCTGGATCGAACTGGGCCCCCACCCTACCACCCTGCCCATGCTGACCGTGCACCCCGGCGTGTCTAAGGAGGCCCTGCTCGTGGGCTCCCTGAAGAAGCGACAAGACGACAACCTGACCCTGTCTTCCTCTCTGTCTCAGCTGTACACCTCTAACGTGCCCGTGCAGTGGCGAGACGTGTTCGCCGACGTGTCTGCCGCCTGTGTGTCTCTGCCCTCTTACCCCTGGCAGAAGTCTAAGTTCTGGGTGGCCTGGAAGGAGGACTCTCCCGCTCCCGCCTCTTCCACCGAGGGCTCTCCCGCCCCCACCAAGGCCTTCAACCCCGTGAACGACTTCGGCATGCTGCAGTCTTGGGCTCAGTTCCCCTCTGCCGCCAACTCTCAGACCGCCATCTTCGAGACTCCCATCTCTCTGCTGAAGACCTCTATCACCGGCCACATCGTGGGCGACGTGCCCCTGTGTCCCGCCTCTGTGTACCACGAGCTGGCCCTGGCCGGCATCGAAGCCTCTAAGGCCCACCTGTCTCTGCCCCTGCAAGGCTCTCACTCTGCCCTGTTCAACATCGACTACGTGAAGCCCCTGGTGTACTCTAAGGACGTGGCCCGAGTGGTGAAGACCACCATCGCCATGAACACCGACGGCTCCGGAACCTTCACCGTGGAGTCTTACGCCGACTCTGAACCCGAGTCTGTGCACTGTTCTGGACAGTTCCGACCCCTGCTGGTGGCCGACACCACTACCAAGTTCAACCGAATGGCCCCCGTGGTGTCTCGACGAACCGCCGCCATCTGTTCTGGCGAGGACGACGAGGCCGAGGTGATCACCACCCGAACCGCCTATGAGATCATCTTCACTCGAGTGGTGCGATACGCCAAGGAGTACCACACCATGAAGAACGTGACTATCTCTAAGAACGGCATGGAGGGCTACGCCATCGTGAAGCTGCCCAAGGACCACGACCGATCTAAGTTCGTGGTGCACCCCGTGTTCATGGACACCATGCTGCACGTGGCCGGCTTCCTGGCCAACATGCAAGGCGGCGACAACGACGCCTACATCTGTTCTAAGGTGAAGTCCGTGAAAGCCGTGCCCTCTCTGATCAACAACGACGCCACTTACGGCGTGTTCGTGGTGAACGCCTGGGTGGAGTCTGAGGGCATCATGCTGTCTGACGCCATCGCCGTGGACATCTCTGAACACGGCCAAATTGTGGCTCAGCTGAAGGGCATGTGTTTCAAGAAGCTGCGACTGAACACCCTGCAGCGATCTCTCGCCATGCATGCCGGCCACACCTCCCCCGCCCCCGCCCCTAAGAGAACCGTGGCCGCCGCTCCCAAGCCCAAGATCACCGAGGTGGCCCCCGCTCTGGGCCCCCGATCTTCTCCCGCCAAGCGATCTGTGGACGTGCAGAACACCGTGCTGAAGATCATCGGCGACACCTGTGGCATCGAGGTGTCTGCCCTCGACGTGAACGCCGACCTGGAGACCTACGGAGTGGACTCTCTCATGTCTATCGAGATCCTGCGAAAGTTCGAGGAGTCTTTCCTGCAGATGCAGTTCGACACCACCATCTTTTCTACCTGTAACAACATCACCGAGCTGGTGCGAGAGATTTCTTCTACCATTGGCTCTCAAGCCGCTACCGCCGTCAACACCCCCGAGACCGCTTCTACTCCCGAGCCTACCCTGCAAGGCGACGCCCCTCAGTCTACCGACGTGCGATCTATCCTGCTGGAGCTGATCTCCTCTTTCACCGGATTCGAGATCTCTTCTTTCGACCTGAACGCCGATGCCGACTCTGCCTACGGCCTGGACAAGTTCCTGTTCATCCCCCTGTTCTCTAAGCTGCAGTCTTTCTTCCCCGACGTGACCCTGGACCCCGCCAAGCCCTCTGTGTGTTCTACCATCGGAGAGCTGCTGGACGAGGTCACCGCCCAAGTGCAAGCCGGCCCTTCCTCTTCCTCTCCCGGCATCGTGGACACCAAGCCCATGTTCGTGTCTGTGCTGGGCCTCGATGAATCTGACATCCAAGACGACACCGAGTTCGAGACCATCGGCCTGGACTCTCTGACCGCCATCGAGGCCCTGCACGCCATTCAGACCAAGTACGGCCTGGAACTGCCCTCTAACCTGTTCGAGCTGCACAACACCGTGAAGGCCGTGAACCAATACATCTCTTCCAAGCAGCCCGGCAAGTCTCCTAAGCCCTCTGAGGAGGCCACCATGGACCCCGACAAGGAGGAGGACCTGTCTGACCTGACCCCCGAGCAAGTGCAGTCTGTGGTGCGAGTGCTGCGACTCGACGAAGTGCCCATGTCTGTGCAGAAGTCTTCCTCTTCTGGCTCTCCCCTGTTCCTGTTCCACGACGGCTCTGGAGCCGTCAACTACCTGCGACGACTGGGCTCTGTGGGCCGAGAGTTCTGGGGCTTCAACAACCCCAACTACGCCACCGGCAAGCCCTGGGGCTCTGTGGAGGCCATGGCCTCTGCCTACGCCGACTACGCCGTGAAGGTGGCCGGCTCTCGACCCGTGATCTTCGGCGGATGGTCTTTCGGAGGCGTGGTGGGCTTCGAGGCCGCCCGACAACTGATGCGACGAGGCGTGCCCGTGAAGGGCGTGGTGCTGATCGACTCTCCCTTCCCCGTGGACCACGTGCCCTCTTCTAACGAGTTCATGGCCGTGACCGCCGGCGCCTTCACCCGAGGCGGCCGAACCCCCATCGGCCGAATGATGTGGAAGCAGCTGCAGCAGAACGCCCCCCTCCTGAAAACCTACGACCCCCGAATCGCCGGCGGCCCCTACCCCCCCCTGGTGCTGCTGCACAACCAAGAGGGCATCCCCCCCGACGCCTTCCTGCCCTACCCCGTGCCCCGATGGATGTCTGAGAAGGGCACCGACCCCTGTCTGCTGGCCGACGACTGGTCTGGCCTGGTGGGCGCCTCTATCAAGGTGATCCACCTGCCCGGCACCCACTTCACCACCTTTGCCACCCCCCACCTCGGCGCCGTGACCCAAGCCCTCGTGGATGGCTGTGCCTACCTGGACGAGCTGTAA
